# Viromics in flat mites from Hawaii shows abundant arrays of viruses, expands the evolutionary origin of plant viruses, and provides a surveillance tool for *Brevipalpus*-transmitted viruses

**DOI:** 10.64898/2026.02.11.705406

**Authors:** Alejandro Olmedo-Velarde, Kota Nakasato, Adriana Larrea-Sarmiento, Michael Melzer

## Abstract

Flat mites (Tenuipalpidae) are diverse phytophagous arthropods, among which *Brevipalpus* species are economically important pests capable of transmitting plant viruses. Brevipalpus-transmitted viruses (BTVs) cause localized infections in plants and are classified into two major groups based on cytopathology and genome organization: BTV-C (genera *Cilevirus* and *Higrevirus*, family *Kitaviridae*) and BTV-N (genus *Dichorhavirus*, family *Rhabdoviridae*). Despite their significance, the virome of tenuipalpid mite vectors remains poorly characterized. Using high-throughput sequencing (HTS), we analyzed virus populations associated with *Brevipalpus* and *Dolichotetranychus* mites collected from multiple plant hosts across two Hawaiian Islands. We identified a diverse assemblage of viral sequences affiliated with *Kitaviridae*, negeviruses, *Picornavirales*, *Narnaviridae*, *Tombusviridae*, *Solemoviridae*, *Ourmiaviridae*, *Reoviridae*, and *Potyviridae*. Near-complete genomes of citrus leprosis virus C2H and hibiscus green spot virus 2 (both BTV-C) were recovered, highlighting the utility of HTS-based viromics for surveillance of BTVs in mite vectors. In addition, multiple divergent virus-like contigs were identified based on viral hallmark genes and sequence divergence, including Brevipalpus-associated negevirus, Brevipalpus-associated bluner-like virus, and Dolichotetranychus-associated cile-like virus, all showing evolutionary affinities to BTV-C-related viruses. Phylogenetic analyses support evolutionary links between negeviruses and kitavirids, consistent with the hypothesis that *Kitaviridae* evolved from arthropod-associated ancestors. While some detected plant viruses may reflect ingestion rather than active replication in mites, this study establishes a robust framework for virome-based surveillance of tenuipalpid mites, advancing our understanding of plant virus evolution and supporting agricultural biosecurity and pest management efforts.

## INTRODUCTION

About 1100 mite species belong to the family Tenuipalpidae, which is currently classified into 38 genera (Beard et al., 2013). Among these, *Brevipalpus* and *Tenuipalpus* together comprise more than 600 described species and represent the most economically important genera within the family. However, only only a subset of tenuipalpid mites are considered major pests, causing direct plant injury or indirect damage through toxin injection or plant virus transmission (Gerson, 2008). The species complexes *B. californicus*, *B. obovatus*, and *B. phoenicis* species are among the most frequently intercepted tenuipalpid mites and include the only confirmed vectors within the family that transmit plant viruses of quarantine concern (Freitas-Astúa et al., 2018; Rodrigues & Childers, 2013). The revision of the *B. phoenicis* species complex identified eight distinct species: *B. ferraguti*, *B. yothersi*, *B. hondurani*, *B. feresi*, *B. phoenicis* sensu stricto (s.s.), *B. tucuman*, *B. papayensis,* and *B. azores* (Beard et al., 2015). To date, virus transmission has been experimentally confirmed for *B. yothersi*, *B. papayensis*, *B. phoenicis* s.s., and *B. californicus* (Kondo et al., 2003; Roy et al., 2015; Ramos-Gonzalez et al., 2017; Nunes et al., 2018), and more recently for *B. azores* (Chabi-Jesus et al., 2023; Olmedo-Velarde et al., 2024). In contrast to *Brevipalpus*, species of *Dolichotetranychus* generally exhibit a more restricted host range and remain comparatively understudied with respect to virus associations and vector potential (Mesa et al, 2009).

*Brevipalpus*-transmitted viruses (BTVs) differ from most plant viruses in that they do not establish systemic infections in their hosts, but instead induce localized lesions at feeding sites (Freitas-Astúa et al., 2018; Rodrigues & Childers, 2013). Based on the cytopathological features observed by transmission electron microscopy, particularly the presence and location of viroplasms, BTVs are classified into two major groups: the cytoplasmic type (BTV-C) and the nuclear type (BTV-N) (Kitajima et al., 2003). These viruses affect economically important crops including citrus, coffee, passion fruit, orchids, and numerous other ornamentals. The localized symptoms caused by these viruses are very similar and characterized by chlorotic or green spots and ringspots that may be present in leaves, fruits, and stems (Freitas-Astúa et al., 2018; Kitajima et al., 2003).

At the taxonomic level, BTV-N belongs to the genus *Dichorhavirus* (*Rhabdoviridae* family), while BTV-C comprises members of the genera *Cilevirus* and *Higrevirus* (*Kitaviridae* family) (Nunes et al., 2017; Freitas-Astua et al., 2018; Olmedo-Velarde et al., 2024). Both BTV groups are phylogenetically related arthropod-infecting viruses: BTV-C shows affinities to negeviruses, whereas BTV-N clusters with rhabdovirids. Although BTV-C and BTV-N differ in genome organization and replication strategies, both cause localized infections in plants, and both are transmitted in a persistent manner (Freitas-Astua et al., 2018). Available evidence supports persistent propagative transmission for BTV-N (Kondo et al., 2025), whereas BTV-C is currently considered circulative and non-propagative in mites, based primarily on ultrastructural studies, with additional molecular and protein-level analyses still required to fully resolve replication dynamics (Tassi et al., 2022). These features have led to the hypothesis that the ancestors of BTV-C and BTV-N colonized arthropods and harbored unsegmented RNA genomes characteristic of negeviruses and classical rhabdovirids, respectively (Kondo et al., 2017; Freitas-Astúa et al., 2018; Ramos-González et al., 2020; Quito-Avila et al., 2020).

In the last decade, virome studies of arthropods have provided valuable insights into virus diversity and the evolutionary relationships between plant- and arthropod-infecting viruses (Shi et al., 2016; Zhang et al., 2018). In this context, negeviruses have been proposed as a diverse group of positive-sense RNA viruses that can be subdivided into several genera-level lineages, including Nelorpivirus, Sandewavirus, Aphiglyvirus, and *Centivirus* (Kallies et al., 2014; Kondo et al., 2020). However, despite the established role of *Brevipalpus* mites as virus vectors, the broader virome associated with tenuipalpid mites, including *Dolichotetranychus*, remains poorly characterized.

In this study, we used high-throughput sequencing (HTS) to characterize BTV-related viruses within the virome associated with *Brevipalpus* and *Dolichotetranychus* mites collected from multiple host plants in Hawaii. Our objectives were to document the diversity of virus-like sequences associated with tenuipalpid mites, recover and characterize BTV genomes where possible, and place newly identified virus-like contigs in an evolutionary context relative to known arthropod- and plant-associated viruses. This study will open new research opportunities to characterize new BTVs found in populations of flat mites from other geographic locations.

## MATERIALS AND METHODS

### Flat mite specimen collection

Leaves, twigs, and/or fruits were collected from several known plant hosts of flat mites (Tenuipalpidae family) (Beard et al. 2013) from several locations across the islands of Oahu and Hawaii, including commercial orchards, city community gardens, and university ornamental gardens (Figure 1). At the time of collection, most plant samples did not display obvious virus-like symptoms, although some were taken from plants that also carried other senescing leaves with confirmed kitavirid symptoms. The plant material was placed in hermetic bags, transported to the Agrosecurity Laboratory at the University of Hawaii at Manoa, and inspected individually under a dissecting microscope. If present, flat mites were retrieved using a sterile fine needle and placed into a 1.5 mL tube containing 500 μL of 95% ethanol. The needle was flame-sterilized between each mite sample collected per plant host per location. Each mite sample consisted of at least ten flat mites collected from the same plant host per location. Capped tubes were securely wrapped with one layer of parafilm and stored at -80°C until nucleic acid extraction was performed. Following the inspection of 35 plant samples representing species in the genera *Ananas*, *Anthurium*, *Areca*, *Carica, Citrus*, *Coffea, Cordyline, Dendrobium*, *Ficus*, *Hibiscus, Oncidium, Passiflora*, *Phalaenopsis*, *Theobroma*, and *Gardenia*, 15 samples were found to be infested with flat mites and were included in this study. Infested host samples included *Citrus* spp. (four samples), *Passiflora edulis* (three samples), *Hibiscus* spp. (three samples), *Coffea* sp. (two samples), *Carica papaya* (two samples), and *Ananas comosus* (one sample) (Figure 1 and Table 1).

**Figure 1.**
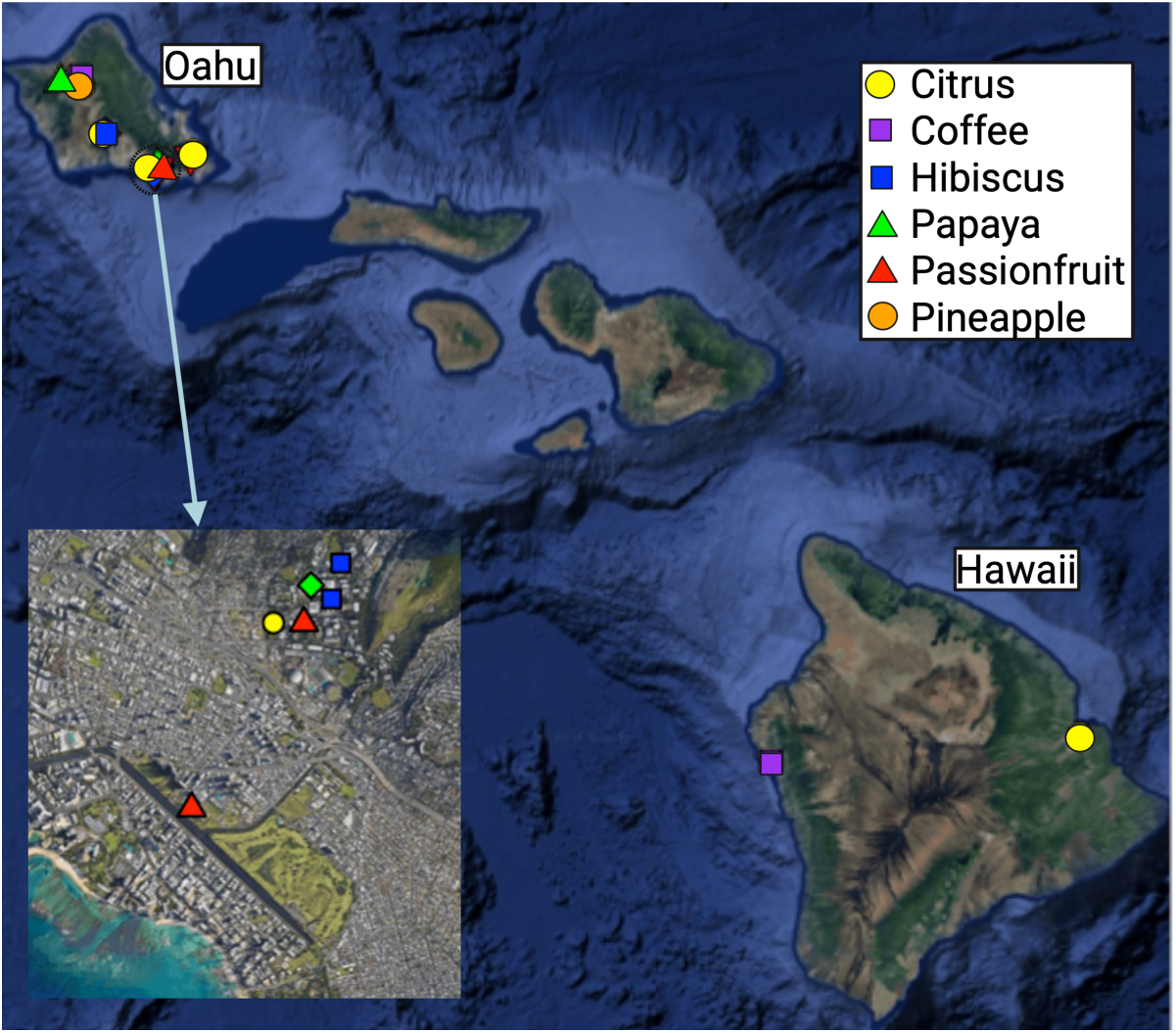
Map detailing growing locations and hosts from which flat mite specimens were collected on the islands of Oahu and Hawaii. One individual mite sample for nucleic acid extraction comprised at least 10 tenuipalpid mites collected per location per plant host.

**Table 1.**
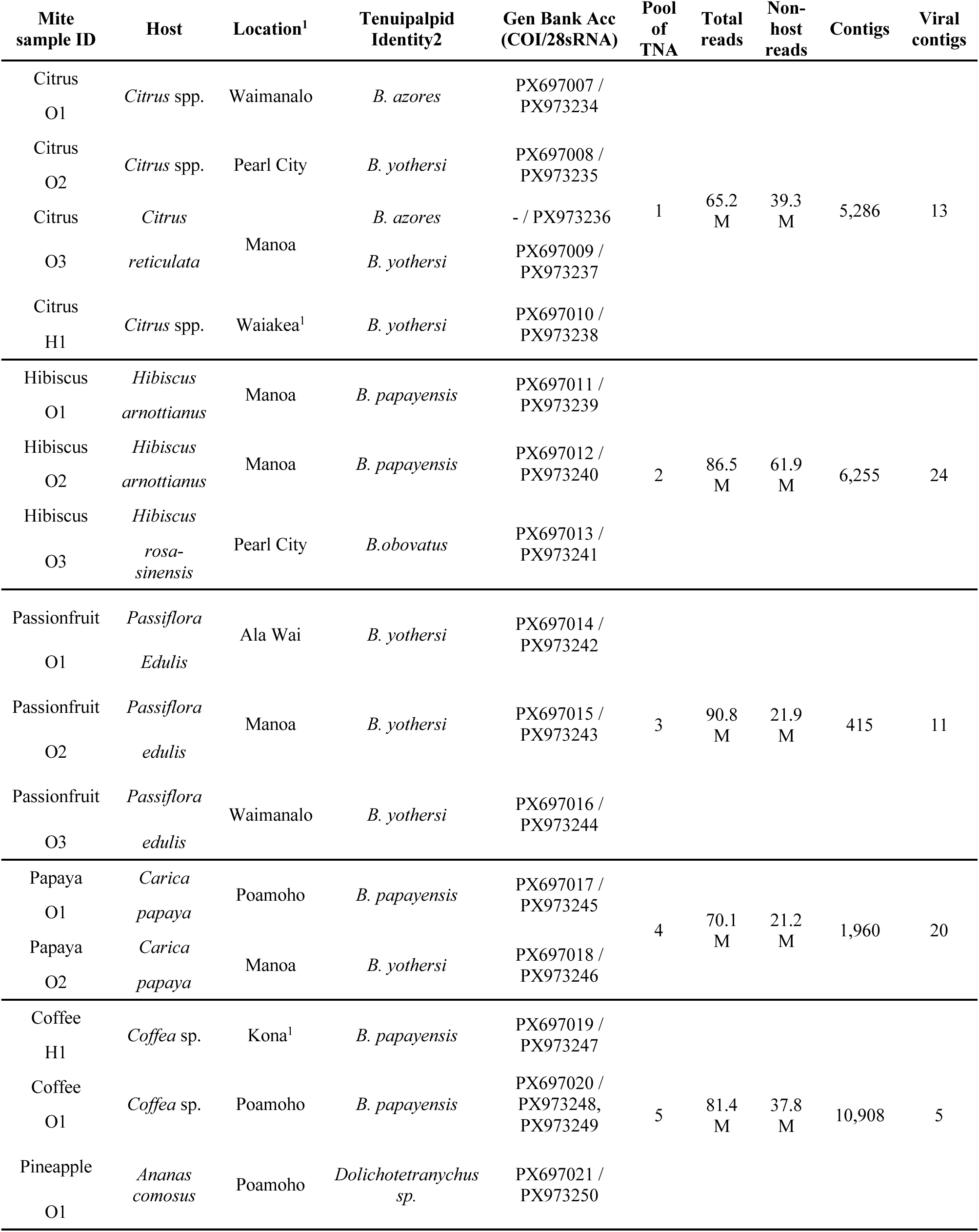
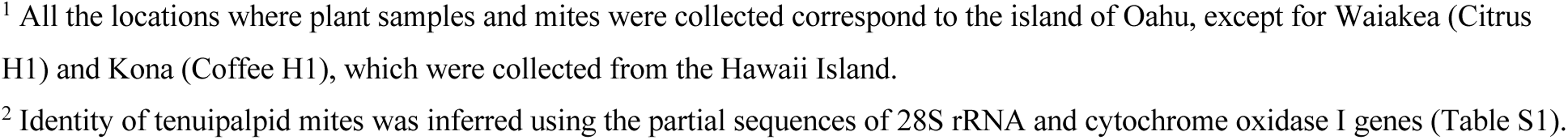
Location, identity of the tenuipalpid mites, number of HTS reads and contigs generated to determine the virome present in each mite sample collected from Oahu and Hawaii Islands.

| Mite sample ID | Host | Location<sup>1</sup> | Tenuipalpid Identity<sup>2</sup> | Gen Bank Acc (COI/28sRNA) | Pool of TNA | Total reads | Non-host reads | Contigs | Viral contigs |
| --- | --- | --- | --- | --- | --- | --- | --- | --- | --- |
| Citrus O1 | *Citrus* spp. | Waimanalo | *B. azores* | PX697007 / PX973234 | 1 | 65.2 M | 39.3 M | 5,286 | 13 |
| Citrus O2 | *Citrus* spp. | Pearl City | *B. yothersi* | PX697008 / PX973235 |  |  |  |  |  |
| Citrus O3 | *Citrus reticulata* | Manoa | *B. azores* | - / PX973236 |  |  |  |  |  |
|  |  |  | *B. yothersi* | PX697009 / PX973237 |  |  |  |  |  |
| Citrus H1 | *Citrus* spp. | Waiakea<sup>1</sup> | *B. yothersi* | PX697010 / PX973238 |  |  |  |  |  |
| Hibiscus O1 | *Hibiscus arnottianus* | Manoa | *B. papayensis* | PX697011 / PX973239 | 2 | 86.5 M | 61.9 M | 6,255 | 24 |
| Hibiscus O2 | *Hibiscus arnottianus* | Manoa | *B. papayensis* | PX697012 / PX973240 |  |  |  |  |  |
| Hibiscus O3 | *Hibiscus rosa-sinensis* | Pearl City | *B. obovatus* | PX697013 / PX973241 |  |  |  |  |  |
| Passionfruit O1 | *Passiflora edulis* | Ala Wai | *B. yothersi* | PX697014 / PX973242 | 3 | 90.8 M | 21.9 M | 415 | 11 |
| Passionfruit O2 | *Passiflora edulis* | Manoa | *B. yothersi* | PX697015 / PX973243 |  |  |  |  |  |
| Passionfruit O3 | *Passiflora edulis* | Waimanalo | *B. yothersi* | PX697016 / PX973244 |  |  |  |  |  |
| Papaya O1 | *Carica papaya* | Poamoho | *B. papayensis* | PX697017 / PX973245 | 4 | 70.1 M | 21.2 M | 1,960 | 20 |
| Papaya O2 | *Carica papaya* | Manoa | *B. yothersi* | PX697018 / PX973246 |  |  |  |  |  |
| Coffee H1 | *Coffea* sp. | Kona<sup>1</sup> | *B. papayensis* | PX697019 / PX973247 | 5 | 81.4 M | 37.8 M | 10,908 | 5 |
| Coffee O1 | *Coffea* sp. | Poamoho | *B. papayensis* | PX697020 / PX973248, PX973249 |  |  |  |  |  |
| Pineapple O1 | *Ananas comosus* | Poamoho | *Dolichotetranychus* sp. | PX697021 / PX973250 |  |  |  |  |  |

<sup>1</sup> All the locations where plant samples and mites were collected correspond to the island of Oahu, except for Waiakea (Citrus H1) and Kona (Coffee H1), which were collected from the Hawaii Island.
<sup>2</sup> Identity of tenuipalpid mites was inferred using the partial sequences of 28S rRNA and cytochrome oxidase I genes (Table S1).

### Nucleic acid extraction

Each mite sample consisted of at least 10 flat mites collected from the same plant host per location. These pooled mite samples (≥10 mites per sample) were used as the minimum input for nucleic acid extraction. Tubes containing the mites were briefly centrifuged for 30 seconds at 2,000xg to pellet the mites, and the ethanol was carefully removed using a micropipette. The mites were then powdered using liquid nitrogen and a sterile micro-pestle, then homogenized with 100 μL of CTAB buffer containing β-mercaptoethanol. Total nucleic acids (TNA) were immediately extracted as described by Li et al. (2008), and the resuspended TNA were further purified using the PureLink RNA mini kit (ThermoFisher Scientific, Waltham, MA), according to the manufacturer’s instructions for liquid samples. TNA was resuspended in 20 µL of DEPC-treated water and quantified using a NanoDrop 2000 Spectrophotometer (ThermoFisher Scientific, Waltham, MA). To reduce sequencing costs, the TNA extracts from each sample, i.e., mites from the same hosts, but different locations, were equimolarly pooled to have five final pool samples (Table 1).

### Determination of the virome in the tenuipalpid mite samples

Pools of TNA were stabilized in RNA Stabilization tubes (Genewiz, South Plainfield, NJ) according to the manufacturer’s instructions. The tubes containing the stabilized TNA pools were sent to the University of California, Irvine Genomics High Throughput Facility. Following the manufacturer’s instructions, ribodepletion and library construction were performed using the SMARTer Stranded Total RNA-Seq Kit v3 - Pico Input Mammalian kit (Clontech, Mountain View, CA). HTS was performed on an Illumina NovaSeq 6000 platform using an S4 200-cycle kit. Genome assembly and bioinformatic analyses were performed as described (Olmedo-Velarde et al., 2019) with slight modifications. Briefly, paired-end reads were trimmed and quality filtered using Trimmomatic 0.35.3 (Bolger et al., 2014).

Tenuipalpid-specific reads were mapped to the draft genome available in GenBank (https://ftp.ncbi.nlm.nih.gov/genomes/genbank/invertebrate/Brevipalpus_yothersi/all_assembly_versions/) and any Tenuipalpidae sequences available in GenBank (last accessed on September 20, 2021) using Bowtie 2 (Langmead et al., 2012). The remaining reads were *de novo* assembled using SPAdes (Bankevich et al., 2012). Contigs were annotated using BLASTX search (Altschul et al., 1997) against the viral genome database (ftp://ftp.ncbi.nih.gov/genomes/Viruses/all.fna.tar.gz) as of November 2021, using an E-value cutoff of 1e−5. Contigs were retained as candidate viral sequences when their best BLASTx hits corresponded to viral proteins and were supported by coherent open reading frame organization, conserved viral domain content when present, and consistent read-mapping support. Contigs similar to virus sequences were then used as a reference for an iterative mapping approach (Dey et al., 2019) using the Geneious mapper plug-in implemented in Geneious v. 10.1.3 (Kearse et al., 2012) and the trimmed sequence reads. To complement viral detection, we also used VirSorter2 (Guo et al., 2021a) to identify viral-like contigs. Kaiju (Menzel et al., 2016) and web BLASTn and BLASTx tools were used to annotate viral-like contigs identified using VirSorter2.

HTS data were deposited in the NCBI Sequence Read Archive (SRA) under BioProject PRJNA1393311 (accessions SAMN54298324–SAMN54298328) and viral contigs were deposited in GenBank under accession PX763617-PX763666.

### Mite species identification by DNA barcoding

Complementary DNA (cDNA) was synthesized using mite TNA samples, random and oligo dT primers, and the Maxima-H minus reverse transcription kit (ThermoFisher Scientific, Waltham, MA) using the manufacturer’s instructions. Two microliters of two-fold diluted cDNA reactions were used in endpoint PCR for DNA barcoding and internal PCR control using the 28S rRNA primers, D1D2w2: 5’-ACAAGTACCDTRAGGGAAAGTTG-3’, 28Sr0990: 5’-CCTTGGTCCGTGTTTCAAGAC-3’ (Sonnenberg et al., 2007; Mironov et al., 2012) that produce an expected amplicon of ∼700 bp. The cytochrome oxidase unit I (COI) gene was additionally amplified using the COI primers, DNF: 5’-TACAGCTCCTATAGATAAAAC-3’, DNR: 5’-TGATTTTTTGGTCACCCAGAAG-3’ (Navajas et al., 1996) that produce an expected amplicon of ∼450bp. All DNA barcoding PCR assays were performed using Q5 High Fidelity DNA Polymerase (New England Biolabs, Ipswich, MA).

### Confirmation of virus presence by RT-PCR assays and Sanger sequencing

Endpoint RT-PCR assays were performed on cDNA synthesized from individual mite samples to confirm the presence of viral sequences detected by HTS. For viral contigs containing conserved domains (such as RdRp, HEL, or MET), primers were designed to target these annotated regions. In cases where no conserved domains were detected, primers were instead designed in regions showing sequence similarity to known viral proteins, as identified by BLASTx. Primer3 (Untergasser et al., 2012) was used for the primer design, considering thermodynamic primer features described previously (Arif & Ochoa-Corona, 2013). These primer sets were used for specific detection of all the Tenuipalpidae-associated viruses in endpoint RT-PCR assays using 0.5 µM as the final primer concentration and 55 °C as the annealing temperature. All barcoding amplicons were cloned into pGEM-T-Easy (Promega, Madison, WI), and 3-5 clones were sequenced. Amplicons from all other RT-PCR assays were gel extracted, purified, and bi-directionally sequenced using the Sanger method. COI sequences were deposited in GenBank under accessions PX697007–PX697021, and 28S rRNA sequences under accessions PX973234-PX973250.

### Virus Genome Sequence Analyses

Virus contigs presenting similarity to BTVs, i.e., viruses belonging to the *Kitaviridae* family or *Dichorhavirus* genus, were selected for further analysis. The NCBI ORFfinder program (www.ncbi.nlm.nih.gov/orffinder) was used to identify putative open reading frames (ORFs) *in silico*, using the standard genetic code and default parameters. Conserved domains were predicted using either the NCBI conserved domain search tool (www.ncbi.nlm.nih.gov/Structure/cdd/wrpsb.cgi) or HMMSCAN (www.ebi.ac.uk/Tools/hmmer/search/hmmscan) implemented in HMMER (Potter et al., 2018).

HMMSCAN was also used for the prediction of transmembrane helices. BLASTP searches were used to retrieve protein homologs and infer the putative function. Pairwise protein sequence comparisons using orthologous sequences retrieved from GenBank were performed using LALIGN (www.ebi.ac.uk/Tools/psa/lalign) (Huang & Miller, 1991). ORFs and putative protein functions were inferred based on sequence homology, conserved domain architecture, and genome organization relative to well-characterized viruses. As is standard in viromics studies, these annotations represent predicted functions derived from strong homology to viral hallmark genes (e.g., RdRp, HEL, MET).

### Phylogenetic Analyses

Phylogenetic relationships were inferred using the predicted protein sequences encoded by contigs presenting similarity to BTVs, and their respective virus homolog sequences available in GenBank. Multiple protein sequence alignments were performed with ClustalW (Thompson et al., 1994) implemented in MEGA 7.0.25 (Kumar et al., 2016) using the RdRp, MET, HEL, and/or coat protein (CP) domains. Ambiguous positions for each alignment were curated using Gblocks 0.91b (https://ngphylogeny.fr) (Talavera & Castresana, 2007) using a minimum block length of four amino acids, allowing gaps in up to half of the sequences, with all other parameters set to default values. The best model of protein evolution (LG+G+I) was used to generate a Maximum Likelihood tree with 1,000 bootstrap repetitions. Similarly, to corroborate the *Brevipalpus* species identity, phylogenetic relationships were inferred using the partial 28S rRNA and COI genes sequences and their respective homolog sequences available in GenBank. Reference sequences were selected primarily from Beard et al. (2015) and related studies that revised the B. phoenicis species complex using integrative taxonomy and detailed morphological validation. Only reference sequences corresponding to species detected in thist stuy, or closely related taxa were included. All reference sequences were aligned over the same homologous regions corresponding to the COI and 28S rRNA fragments amplified using the primers described above, ensuring direct compatibility across datasets. Alignment curation, and phylogenetic tree construction, using GTR+G+I as the best model of DNA evolution, were performed as detailed above.

### Virus nomenclature

Viral contigs were classified into putative taxa based on a combination of: (i) BLASTx similarity to viral hallmark domains (RdRp, MET, HEL), (ii) amino acid identity thresholds (<70% aa identity in hallmark domains considered distinct), (iii) nucleotide similarity (<70% nt identity by BLASTn considered distinct), (iv) predicted ORF organization, and (v) contig-specific read mapping patterns. Contigs within the same viral family/order (e.g., Picornavirales, Reoviridae, Kitaviridae, Negevirus-like) were compared across these metrics to determine whether they represented independent genomic fragments or segments of a single viral genome. Multi-segment viruses (e.g., Brevipalpus-associated reovirus RNAs 1–12, Brevipalpus-associated bluner-like virus RNAs 1–2, and CiLV-C2 and HGSV-2 RNAs 1–3) were defined when contigs showed complementary domain content, compatible genome organization, and overlapping presence/coverage within the same library. By contrast, contigs within the same group that showed low aa/nt identity in overlapping regions and distinct ORF organization and/or coverage patterns (e.g., Brevipalpus-associated picornavirus 1–13, ourmiavirus 1–5, narnavirus 1–3, tombusvirus 1–3) were treated as separate candidate viral species. In total, 41 virus-derived contigs that did not correspond to known viruses were grouped into 30 putative new viruses (Table 2), including partial virus-like contigs from Dolichotetranychus mites (Dolichotetranychus-associated negevirus contig and Dolichotetranychus-associated cile-like virus contig). The virus contigs identified in this study have been named using the following nomenclature system: (i) the first part of the name is the genus of the mite associated with the source of the virus; (ii) the second part identifies the virus taxonomic group (order, family, or genus) or virus-like lineage to which the contig was most closely related. This nomenclature system was adopted solely for this study and does not reflect official virus names or formal proposals to the International Committee on Taxonomy of Viruses; all entities described here should be regarded as candidate viruses requiring further confirmation (e.g., recovery of full genomes and biological characterization).

**Table 2.** Putative novel viruses identified in *Brevipalpus* and *Dolichotetranychus* mites from Hawaii, highlighting their tentative taxonomic affiliations and relationships with their closest homologs.

| *Brevipalpus*-associated Virome |  |  |  |  |  |
| --- | --- | --- | --- | --- | --- |
| Citrus |  |  |  |  |  |
| New Virus Name | GenBank Accession | Contig Size (bp) | FPKM | Closest Hit (Accession; length in bp) | Identity* (Query Coverage) |
| Brevipalpus-associated ourmiavirus 1 | PX763618 | 1238 | 2.9 | Erysiphales ourmia-like virus 1 (MN628265; 2190) | 44.4% (90%) |
| Brevipalpus-associated narnavirus 1 | PX763617 | 2185 | 52.8 | Plasmopara viticola lesion associated virus 22 (MN539839; 2327) | 40% (94%) |
| Brevipalpus-associated picornavirus 1 | PX763619 | 9016 | 8.7 | Hubei picorna-like virus 78 (NC_033212; 9440) | 39.6% (58%) |
| Brevipalpus-associated picornavirus 2 | PX763620 | 8923 | 125.4 | Eotetranychus kankitus picorna-like virus (MN745092; 9772) | 51.8% (62%) |
| Brevipalpus-associated picornavirus 3 | PX763621 | 6719 | 2.4 | Darwin bee virus 4 (MG995699; 8838) | 38% (67%) |
| Brevipalpus-associated picornavirus 4 | PX763622 | 2179 | 1.9 | Budalangi Iflavi-like virus (MZ078286; 9758) | 29.4% (80%) |
| Brevipalpus-associated picornavirus 5 | PX763623 | 1551 | 1.1 | Bundaberg bee virus 4 (MG995705; 8339) | 49.6% (99%) |
| Brevipalpus-associated picornavirus 6 | PX763624 | 1044 | 1.7 | Bundaberg bee virus 4 (MG995705; 8339) | 43.1% (83%) |
| Brevipalpus-associated tobusvirus 1 | PX763625 | 2298 | 3.0 | Erysiphales associated ambiguivirus 1 (MN609869; 3350) | 51.7% (61%) |
| Brevipalpus-associated tobusvirus 2 | PX763626 | 1301 | 1.3 | Macrophomina phaseolina umbra-like virus 1 (MT062435; 3796) | 46.6% (55%) |
| Hibiscus |  |  |  |  |  |
| New Virus Name | GenBank Accession | Contig Size (bp) | FPKM | Closest Hit (Accession; length in bp) | Identity* (Query Coverage) |
| Brevipalpus-associated ourmiavirus 2 | PX763632 | 2497 | 2.1 | Magnaporthe oryzae ourmia-like virus 5 (MW752173; 2792) | 45.3% (74%) |
| Brevipalpus-associated narnavirus 2 | PX763629 | 2422 | 6.1 | Erysiphe necator associated narnavirus 21 (MN605434; 2484) | 38.8% (96%) |
| Brevipalpus-associated ourmiavirus 3 | PX763633 | 2279 | 4.8 | Erysiphe necator associated ourmia-like virus 115 (MN611642; 2596) | 51.1% (73%) |
| Brevipalpus-associated ourmiavirus 4 | PX763634 | 2250 | 6.5 | Erysiphe necator associated ourmia-like virus 7 (MN552396; 2250) | 43.9% (80%) |
| Brevipalpus-associated narnavirus 3 | PX763630 | 2233 | 1.6 | Plasmopara viticola lesion associated narnavirus 22 (MN539839; 2327) | 40.1% (96%) |
| Brevipalpus-associated ourmiavirus 5 | PX763635 | 2216 | 3.8 | Magnaporthe oryzae botourmiavirus 2 (MW752174; 2260) | 41.9% (75%) |
| Brevipalpus-associated picornavirus 7 | PX763636 | 9238 | 2.9 | Sacbrood virus (NC_002066; 8832) | 29.9% (76%) |
| Brevipalpus-associated picornavirus 8 | PX763637 | 6168 | 1.6 | Leuven Picorna-like virus 1 (MZ443589; 9022) | 28.3 (80%)% |
| Brevipalpus-associated picornavirus 9 | PX763638 | 4707 | 19.2 | Aphis glycines virus 2 (NC_028381; 4850) | 33.7% (58%) |
| Brevipalpus-associated picornavirus 10 | PX763639 | 2700 | 2.7 | La Jolla virus (MH384324; 10250) | 31.6% (80%) |
| Brevipalpus-associated picornavirus 11 | PX763640 | 1246 | 0.6 | Xysticus cristatus iflavirus (KX898495; 9274) | 44.5% (39%) |
| Brevipalpus-associated tombusvirus 3 | PX763641 | 3801 | 1.2 | Erysiphales associated ambiguvirus 1 (MN609869; 3350) | 60.1% (37%) |
| Brevipalpus-associated negevirus | PX763631 | 9683 | 2.9 | Fort Crockett virus (NC_034157; 7903) | 41.3% (55%) |
| Brevipalpus-associated bluner-like virus RNA 1 | PX763627 | 3699 | 2.4 | Erysiphe necator associated bluner-like virus 1 (MN630180; 3581) | 42.7% (91%) |
| Brevipalpus-associated bluner-like virus RNA 2 | PX763628 | 3549 | 3.6 | Hubei virga-like virus 17 (NC_033222; 9481) | 31.6% (26%) |
| Hibiscus green spot virus 2 RNA 1 | PX763642 | 8222 | 1.7 | Hibiscus green spot virus 2 (NC_016141; 8372) | 98.9% (97%) |
| Hibiscus green spot virus 2 RNA 2 | PX763643 | 2954 | 1.4 | Hibiscus green spot virus 2 (NC_016143; 3169) | 99.1% (35%) |
| Hibiscus green spot virus 2 RNA 3 | PX763644 | 3128 | 3.8 | Hibiscus green spot virus 2 (NC_016142; 3113) | 98.7% (28%) |

| Brevipalpus-associated ourmiavirus 6 contig |  | 2338 | - | Sjack associated botourmia-like virus 29<br>(OQ987314.1; 2334) | 76.69%<br>(76%) |
| --- | --- | --- | --- | --- | --- |
| Brevipalpus-associated ourmiavirus 7 contig |  | 1882 | - | Mbeech associated botourmia-like virus 53<br>(OQ987288.1; 2201) | 65.79%<br>(97%) |
| **Passionfruit** |  |  |  |  |  |
| **New Virus Name** | **GenBank Accession** | **Contig Size (bp)** | **FPKM** | **Closest Hit (Accession; length in bp)** | **Identity\* (Query Coverage)** |
| Brevipalpus-associated reovirus RNA 1 | PX763650 | 3962 | 7.0 | Diaphorina citri cimodo-like virus (MZ484733; 4084) | 28.5%<br>(63%) |
| Brevipalpus-associated reovirus RNA 2 | PX763651 | 3917 | 15.3 | Diaphorina citri cimodo-like virus (MZ484734; 3713) | 22.9%<br>(44%) |
| Brevipalpus-associated reovirus RNA 3 | PX763652 | 3651 | 27.2 | Shenzhen reo-like virus 5 (MW177754; 3307) | 27.9%<br>(13%) |
| Brevipalpus-associated reovirus RNA 4 | PX763653 | 2146 | 27.3 | Diaphorina citri cimodo-like virus (MZ484736; 2306) | 23.3%<br>(82%) |
| Brevipalpus-associated reovirus RNA 5 | PX763654 | 2138 | 1.3 | Hubei reo-like virus 2 (KX884662; 4128) | 30.3%<br>(83%) |
| Brevipalpus-associated reovirus RNA 6 | PX763655 | 1819 | 61.0 | Cimodo virus (NC_024920; 1874) | 24%<br>(88%) |
| Brevipalpus-associated reovirus RNA 7 | PX763656 | 1828 | 0.7 | Diaphorina citri cimodo-like virus (MZ484733; 4084) | 34.2%<br>(31%) |
| Brevipalpus-associated reovirus RNA 8 | PX763657 | 1709 | 1.4 | Diaphorina citri cimodo-like virus (MZ484734; 3713) | 24.7<br>(69%)% |
| Brevipalpus-associated reovirus RNA 9 | PX763658 | 1416 | 1.3 | Diaphorina citri cimodo-like virus (MZ484736; 2306) | 27.1%<br>(85%) |
| Brevipalpus-associated reovirus RNA 10 | PX763659 | 1235 | 0.6 | Cimodo virus (NC_024920; 1874) | 23.3%<br>(38%) |
| Brevipalpus-associated reovirus RNA 11 | PX763660 | 845 | 0.6 | French Guiana reovirus (MT792646; 2585) | 23.4%<br>(70%) |
| Brevipalpus-associated reovirus RNA 12 | PX763661 | 727 | 0.5 | Cimodo virus (NC_024920; 1874) | 31.9%<br>(87%) |
| Citrus leprosis virus C2 RNA 1 | PX763662 | 8740 | 102.0 | Citrus leprosis virus C2 (MW413437; 8536) | 99.6%<br>(87%) |
| Citrus leprosis virus C2 RNA 2 | PX763663 | 5052 | 119.0 | Citrus leprosis virus C2 (MW413438 ; 4878) | 98.5%<br>(34%) |
| **Papaya** |  |  |  |  |  |
| **New Virus Name** | **GenBank Accession** | **Contig Size (bp)** | **FPKM** | **Closest Hit (Accession; length in bp)** | **Identity\* (Query Coverage)** |

| Brevipalpus-associated<br>picornavirus 12 | PX763645 | 1771 | 1.2 | Bemisia tabaci picorna-like<br>virus 1<br>(MW256668; 10084) | 32.8%<br>(98%) |
| --- | --- | --- | --- | --- | --- |
| Brevipalpus-associated<br>picornavirus 13 | PX763646 | 2707 | 0.7 | Diaphorina citri picorna-like<br>virus (KT698837; 8496) | 35%<br>(65%) |
| Citrus leprosis virus<br>C2 RNA 1 | PX763647 | 8676 | 7.4 | Citrus leprosis virus C2<br>(MW413437; 8536) | 99.6%<br>(87%) |
| Citrus leprosis virus<br>C2 RNA 2 | PX763648 | 4801 | 6.5 | Citrus leprosis virus C2<br>(MW413438 ; 4878) | 98.5%<br>(34%) |
| Papaya ringspot<br>virus | PX763649 | 3686 | 0.5 | Papaya ringspot virus<br>(NC_001785; 10326) | 96.5%<br>(100%) |
| ***Dolichotetranychus*-associated Virome** |  |  |  |  |  |
| **Pineapple** |  |  |  |  |  |
| **New Virus**<br>**Name** | **GenBank**<br>**Accession** | **Contig**<br>**Size**<br>**(bp)** | **FPKM** | **Closest Hit**<br>**(Accession; length in bp)** | **Identity\***<br>**(Query**<br>**Coverage)** |
| Dolichotetranychus-associated negevirus<br>contig | PX763666 | 2089 | 1.0 | Hubei virga-like virus 4<br>(KX883814; 7920) | 32.8%<br>(98%) |
| Dolichotetranychus-associated solemovirus | PX763664 | 1440 | 2.9 | Neohydatothrips associated<br>sobemo-like virus 1<br>(MN714681; 1555) | 31.8%<br>(24%) |
| Dolichotetranychus-associated cile-like<br>virus contig | PX763665 | 2147 | 1.5 | Hibiscus yellow blotch virus<br>(MT472637; 8382) | 50.6%<br>(89%) |

\* Protein identities were obtained through BLASTx searches using virus contig sequences as query. Due to the nucleotide-protein comparison through BLASTx, some virus contigs may present low query coverage.

### Diversity Measurement

Viral community diversity was analyzed using the vegan package (Oksanen et al., 2025) in R (v. 4.4.2). Two complementary datasets were used: (1) the number of viral species detected within each assigned taxonomic group (species-count dataset) and (2) counts per million reads (CPM), which is normalized abundance values associated with each taxonomic group (abundance dataset). For both datasets, α-diversity (within-sample diversity) was calculated using species richness, Shannon, and Pielou’s evenness indices to describe the complexity and evenness of viral communities in each pool. β-diversity (between-sample dissimilarity) was assessed using Jaccard distances for presence/absence data and Bray–Curtis dissimilarities for abundance data. To aid interpretation, pairwise comparisons are reported as similarity values (derived as 1 – dissimilarity). Non-metric multidimensional scaling (NMDS) was performed with the metaMDS function (k = 2, trymax = 100) to visualize overall community similarity among pools. Because the dataset comprised only five pooled samples, the ordination yielded a near-zero stress value, indicating a trivial fit. Accordingly, NMDS results are interpreted qualitatively as exploratory visualizations of compositional differences rather than quantitative inferences of community structure.

## RESULTS

### Molecular identification of mite samples

Sanger sequencing of the partial sequences of 28S rRNA and COI genes identified the presence of *B. yothersi*, *B. papayensis*, *B. azores,* and *B. obovatus* infesting the collected citrus, hibiscus, coffee, and papaya samples (Tables 1 and S1). In contrast, the flat mites collected from pineapple were identified as *Dolichotetranychus* sp (Tenuipalpidae family). Interestingly, in citrus O3, 28S rRNA sequences highly identical to both *B. azores* and *B. yothersi* were recovered, indicating co-occurrence of these two species on the same host plant. In addition, two closely related *B. papayensis* 28S rRNA genotypes sharing 98.8% nucleotide identity were detected. All the sequences obtained from the barcoding assays presented >98% nucleotide identity to their closest GenBank accession (Table S1).

### Analysis of the virome present in tenuipalpid mites

The five libraries (pools 1-5) generated for this study were composed of mites from four citrus-associated samples (pool 1), three hibiscus-associated samples (pool 2), three passionfruit-associated samples (pool 3), two papaya-associated samples (pool 4), and two coffee-associated and one pineapple-associated sample (pool 5). Illumina HTS generated from ∼65.2 to ∼90.8 M of 100 bp paired-end reads from each library. After trimming, quality control, and subtraction of reads of tenuipalpid origin, a variable number between ∼21.2 M to ∼61.9 M of paired-end reads for the five libraries were *de novo* assembled. The SPAdes *de novo* assembler produced 415 to 10,908 contigs for the five library datasets. BLASTx searches using all the generated contigs from the five library datasets suggested between 5 to 24 viral contigs from several distinct taxa in the five datasets (Table 1). Viral sequences similar to those within *Picornavirales* and the family *Kitaviridae* comprised most of the viral sequences identified in the tenuipalpid specimens. In comparison, a smaller number of contigs were predicted to have similarities with viruses within the families *Narnaviridae*, *Ourmiaviridae*, *Reoviridae*, *Tombusviridae*, *Solemoviridae,* and the unofficial Negevirus taxon (Figure 2A).

**Figure 2.**
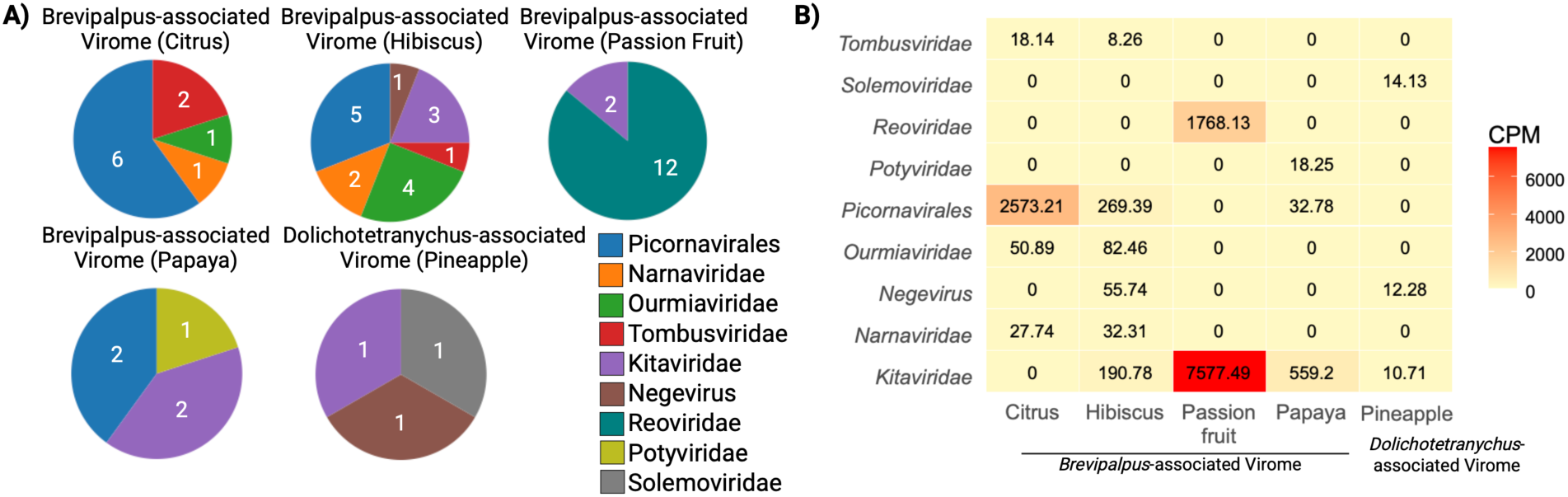
Composition and abundance of viruses identified in *Brevipalpus* and *Dolichotetranychus* mites across citrus, hibiscus, papaya, passion fruit, and pineapple. A) Pie charts showing relative abundance and diversity of virus families, orders, or taxons. B) Heatmap illustrating relative read counts (CPM: count per million bases) associated each virus family, order or taxon.

The contigs of putative viral origin identified across the five datasets ranged in length from ∼1 to ∼9 kbp. Most nucleotide sequences likely represented putative new virus species, based on low protein identities (<60%) to their GenBank counterparts (Table 2). Exceptions included contigs showing >96% protein identity to the hibiscus strain of citrus leprosis virus C2 (CiLV-C2H, KC626783-4), hibiscus green spot virus 2 (HGSV-2, HQ852052-4), both belonging to the *Kitaviridae* family, and papaya ringspot virus (PRSV, MT470188), which belongs to the *Potyvirus* genus (*Potyviridae* family). Some contigs likely represent near-complete genome sequences when compared to the genome size and organization of their closest relative (Table 2).

Mapping of the trimmed reads against the putative virus sequences corroborated that they were distributed through the entire sequence, with fragments per kilobase of transcript per million mapped reads (FPKM) values ranging from 0.5 to 125.4 (Table 2), corresponding to less than 0.5% of the total reads for each dataset (Figure 2B).

The relative contribution of viral taxa differed among host-associated pools. For the citrus-derived pool, almost the totality of the viral reads (96%) mapped to contig sequences putatively classified within the order *Picornavirales*, while the remaining reads mapped to contigs classified within the *Narnaviridae* and *Tombusviridae* families (Figure 2B). For the hibiscus-derived pool, about 42%, 30%, and 18% of viral reads mapped to contigs that presented similarity to viruses within the order *Picornavirales*, *Narnaviridae* family, and hibiscus green spot virus 2 (*Kitaviridae* family), respectively. The remaining reads mapped to virus contig sequences putatively classified within the Negevirus taxon (9%) and *Tombusviridae* family (1%). In both pools derived from passionfruit and papaya, the majority of reads mapped to virus contigs that shared >98% nucleotide identity with the cilevirus CiLVC-C2H (*Kitaviridae* family; KC626783-4) (Figure 2B). In addition, for passionfruit- and papaya-derived pools, about 19% and 3% of the viral reads mapped to virus contigs within the *Reoviridae* and *Potyviridae* families, respectively. Although the number of viral reads mapped to the virus contig sequences of potyvirid origin was low (387, Figure 2B), they mapped to contig sequences distributed along the potyvirid homolog genome (data not shown).

These contig sequences presented ∼96% nucleotide identities to PRSV isolates from Taiwan (MT470188) and Hawaii (EU126128). A small number of reads (695) mapped to virus contigs presented similarity to viruses within the order *Picornavirales* in the papaya-derived pool. Finally, for coffee/pineapple-derived pool, the number of viral reads was almost equally mapped in number to virus contig sequences within the *Sobemoviridae* and *Kitaviridae* families and the unofficial taxon Negevirus (Figure 2).

VirSorter2 was able to detect 20 out of 50 viral contigs that were identified through our BLASTx-based identification and also detected two putative ourmiavirus-like contigs from Hibiscus-associated samples, that were not identified using the BLASTx approach (Table2). These additional contigs were discovered after original biological materials were no longer available. As such, critical validation through RT-PCR was not completed and these viral contigs were not included in the summary figures and analysis.

### Confirmation of virus presence in individual mite samples

Contig-specific primers were designed in conserved virus domains (RdRp, Met or HEL) or the regions where the nucleotide sequences presented protein homology to the virus sequences in GenBank (Table S2). These primers were used in RT-PCR assays to validate HTS results and virus presence. Most virus contigs were detected in at least one mite sample, except for the *Brevipalpus*-associated narnavirus 2 and *Brevipalpus*-associated picornavirus 3 and 4 from citrus, which were not detected in any of the citrus mite samples. *Brevipalpus*-associated picornavirus contigs 1, 5, and 6 from citrus, and CiLV-C2H RNA 1 and 2 from passionfruit were detected in two individual mite samples. CiLV-C2H RNA 1 and 2 were detected in the mite sample papaya O2. Both papaya O2 and passionfruit O2 mite samples were collected from plants growing in the same garden on the UH Manoa campus. Overall, mite samples citrus O1 and O3, hibiscus O1 and O3, passionfruit O1, O2, and O3, papaya O1 and O2, and pineapple O1 tested positive for at least one virus. In contrast, no virus contigs were detected in the other mite samples. (Table 3). Direct sequencing of all the amplicons corroborated the virus identity by showing >99% nucleotide identity to the original contig sequences. Notably, although three viral contigs were initially detected in the pooled coffee–pineapple TNA library, RT-PCR assays showed that all of these viruses originated exclusively from the pineapple mite sample (Pineapple O1), with neither of the two coffee-derived mite samples testing positive. Considering the scope of this study, virus contig sequences presenting similarity to members classified within the *Kitaviridae* family and the Negevirus taxon underwent further analysis.

**Table 3.**
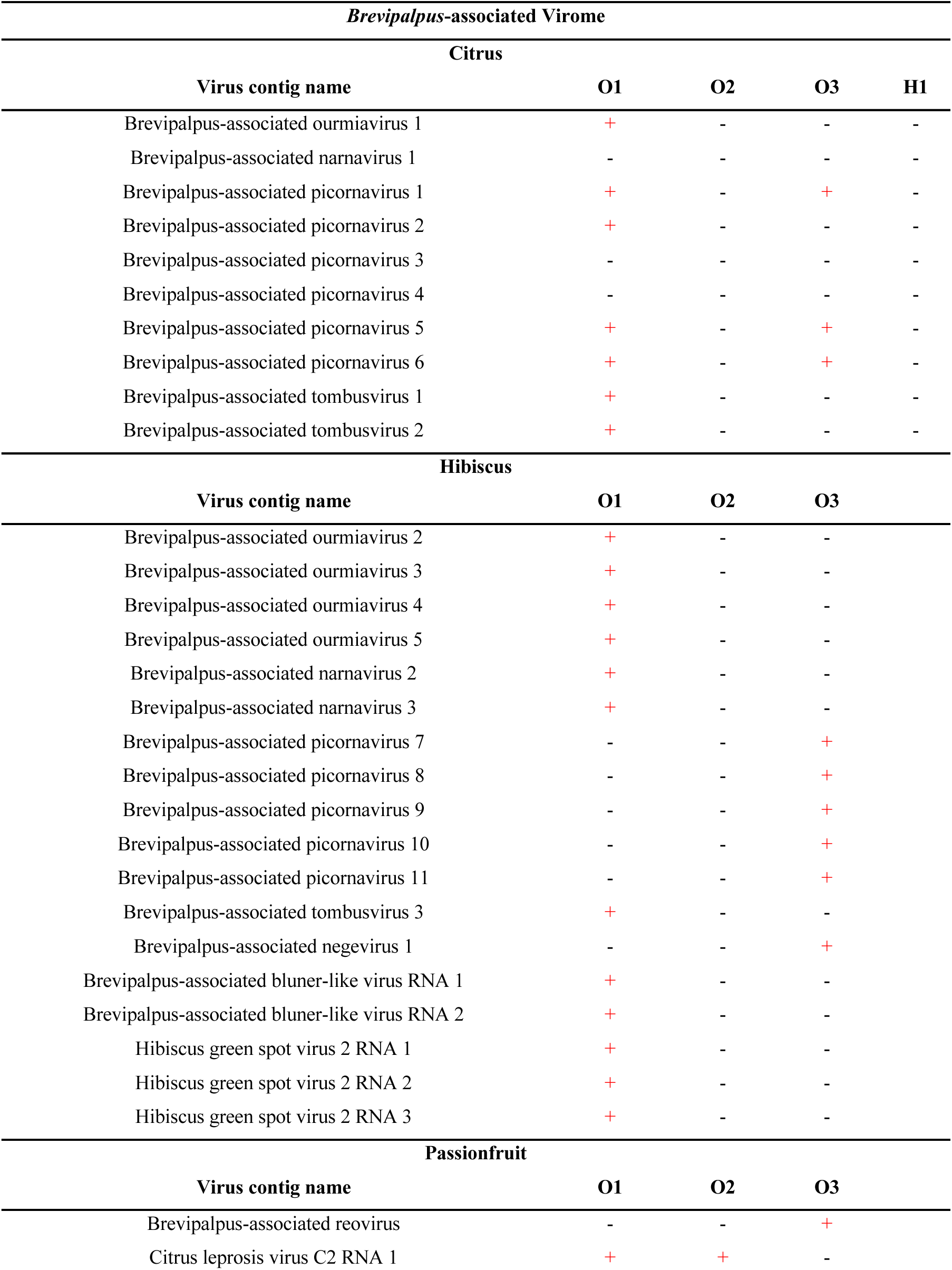
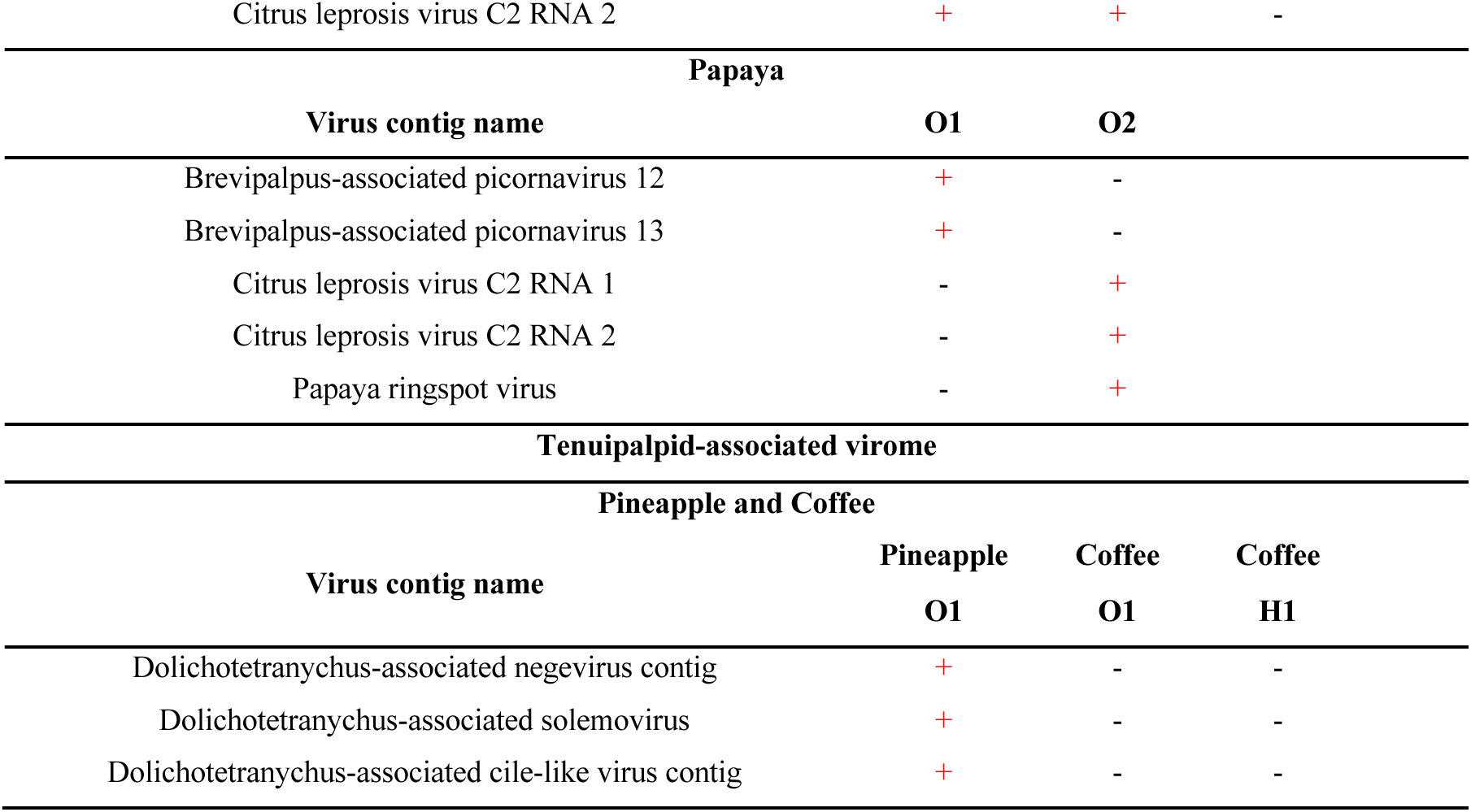
RT-PCR assays results for virus presence confirmation. Virus contig sequences were *de novo* assembled using SPAdes and their viral origin was annotated using BLASTx searches.

| *Brevipalpus*-associated Virome |  |  |  |  |
| --- | --- | --- | --- | --- |
| Citrus |  |  |  |  |
| Virus contig name | O1 | O2 | O3 | H1 |
| Brevipalpus-associated ourmiavirus 1 | + | - | - | - |
| Brevipalpus-associated narnavirus 1 | - | - | - | - |
| Brevipalpus-associated picornavirus 1 | + | - | + | - |
| Brevipalpus-associated picornavirus 2 | + | - | - | - |
| Brevipalpus-associated picornavirus 3 | - | - | - | - |
| Brevipalpus-associated picornavirus 4 | - | - | - | - |
| Brevipalpus-associated picornavirus 5 | + | - | + | - |
| Brevipalpus-associated picornavirus 6 | + | - | + | - |
| Brevipalpus-associated tombusvirus 1 | + | - | - | - |
| Brevipalpus-associated tombusvirus 2 | + | - | - | - |
| Hibiscus |  |  |  |  |
| Virus contig name | O1 | O2 | O3 |  |
| Brevipalpus-associated ourmiavirus 2 | + | - | - |  |
| Brevipalpus-associated ourmiavirus 3 | + | - | - |  |
| Brevipalpus-associated ourmiavirus 4 | + | - | - |  |
| Brevipalpus-associated ourmiavirus 5 | + | - | - |  |
| Brevipalpus-associated narnavirus 2 | + | - | - |  |
| Brevipalpus-associated narnavirus 3 | + | - | - |  |
| Brevipalpus-associated picornavirus 7 | - | - | + |  |
| Brevipalpus-associated picornavirus 8 | - | - | + |  |
| Brevipalpus-associated picornavirus 9 | - | - | + |  |
| Brevipalpus-associated picornavirus 10 | - | - | + |  |
| Brevipalpus-associated picornavirus 11 | - | - | + |  |
| Brevipalpus-associated tombusvirus 3 | + | - | - |  |
| Brevipalpus-associated negevirus 1 | - | - | + |  |
| Brevipalpus-associated bluner-like virus RNA 1 | + | - | - |  |
| Brevipalpus-associated bluner-like virus RNA 2 | + | - | - |  |
| Hibiscus green spot virus 2 RNA 1 | + | - | - |  |
| Hibiscus green spot virus 2 RNA 2 | + | - | - |  |
| Hibiscus green spot virus 2 RNA 3 | + | - | - |  |
| Passionfruit |  |  |  |  |
| Virus contig name | O1 | O2 | O3 |  |
| Brevipalpus-associated reovirus | - | - | + |  |
| Citrus leprosis virus C2 RNA 1 | + | + | - |  |

|  |  |  |  |
| --- | --- | --- | --- |
| Citrus leprosis virus C2 RNA 2 | + | + | - |
| **Papaya** |  |  |  |
| **Virus contig name** | **O1** | **O2** |  |
| Brevipalpus-associated picornavirus 12 | + | - |  |
| Brevipalpus-associated picornavirus 13 | + | - |  |
| Citrus leprosis virus C2 RNA 1 | - | + |  |
| Citrus leprosis virus C2 RNA 2 | - | + |  |
| Papaya ringspot virus | - | + |  |
| **Tenuipalpid-associated virome** |  |  |  |
| **Pineapple and Coffee** |  |  |  |
| **Virus contig name** | **Pineapple**<br>**O1** | **Coffee**<br>**O1** | **Coffee**<br>**H1** |
| Dolichotetranychus-associated negevirus contig | + | - | - |
| Dolichotetranychus-associated solemovirus | + | - | - |
| Dolichotetranychus-associated cile-like virus contig | + | - | - |

### Molecular characterization of virus contig sequences similar to kitavirids and negeviruses

#### Citrus leprosis virus C2H and hibiscus green spot virus 2

Near-complete genomes of the kitavirids CiLV-C2H and HGSV-2 were *de novo* assembled using SPAdes from the datasets originating from passionfruit and papaya, and hibiscus mite datasets, respectively. RT-PCR assays confirmed the presence of CiLV-C2H in the mite samples passionfruit O1, O2, papaya O2, and HGSV-2 in hibiscus O1 (Table 3). The near-complete genomes of CiLV-C2H RNA 1 and 2 that were retrieved from the passionfruit and papaya datasets presented ∼99.9% nucleotide identity between each other following pairwise alignments. Both genomic sequences of CiLV-C2H from the passionfruit and papaya mite datasets presented >97% nucleotide identity to CiLV-C2H previously characterized from hibiscus (KC626783-KC626784). Moreover, the near-complete genome of HGSV-2 from the hibiscus mite dataset presented >98.8% nucleotide identity to HGSV-2 characterized from *C. volkameriana* (HQ852052-HQ852054). The genomic organizations of both CiLV-C2H and HGSV-2 virus sequences were identical to those previously reported (Melzer et al., 2013; Olmedo-Velarde et al., 2024) (Figure 3).

**Figure 3.**
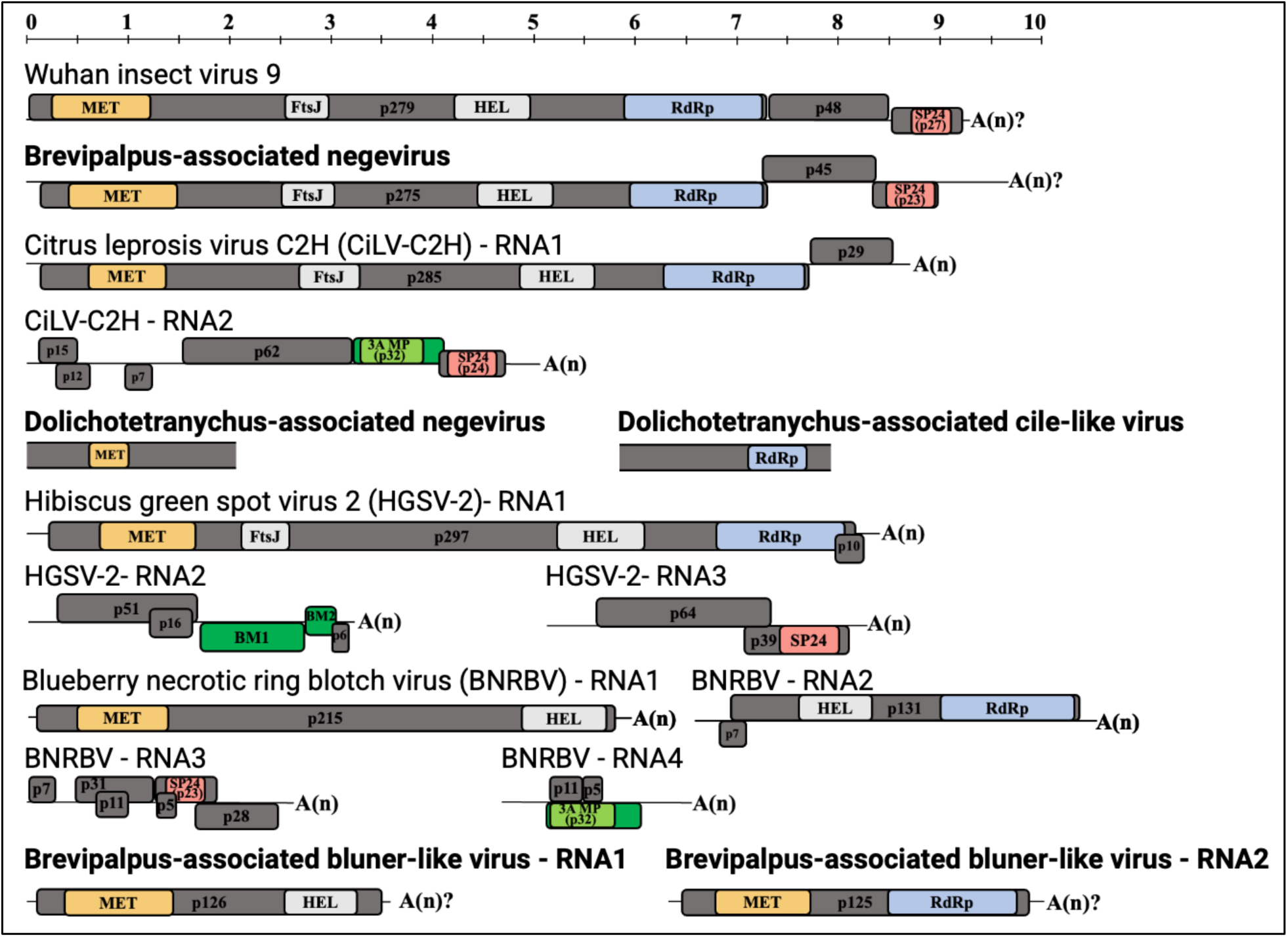
Genome organization of the BTVs-C, citrus leprosis virus C2 (CiLV-C2), hibiscus green spot virus 2 (HGSV-2), and the putative new viruses (bold) that show relationship to members of the *Kitaviridae* family and the unofficial Negevirus taxon. The genomes of Wuhan insect virus 9 (unclassified virus within the Negevirus taxon) and blueberry necrotic ring blotch virus (BNRBV, Genus *Blunervirus, Kitaviridae* family) were included in this figure for genome organization comparisons. Gray and green colored boxes represent open reading frames (ORFs). Green-colored boxes represent ORFs coding for putative genes associated with virus movement. Small ORFs shorter than 300 bp that code for putative proteins containing transmembrane domains are also depicted. Protein conserved domains predicted based on sequence homology and HMM-based searches are represented by colored boxes inside ORFs: methyltransferase (MET, light orange and FtsJ, light gray), RNA-dependent RNA polymerase (RdRp, light blue), helicase (HEL, light gray), movement protein 3A (3A MP, light green), and virion membrane protein of plant and insect viruses (SP24, light red). Based on the confirmed existence of poly-A tails [A(n)] in several negeviruses and kitavirids, it is hypothesized the same feature is present in the putative new viruses [A(n)?]. Genomes were drawn to scale.

#### Contig sequences representing putative new viruses related to kitavirids

Using BLASTx searches, inferred protein sequences from one contig (2,147 bp) found in the pineapple-coffee-derived pool presented ∼51% identity to the kitavirid hibiscus yellow blotch virus (HYBV) (Olmedo-Velarde et al., 2021). Despite the fragmented ORF structure observed at the nucleotide level, a conserved RdRp domain spanning multiple coding regions was detected. For descriptive and comparative purposes, a single continuous ORF was inferred to represent this contig, acknowledging potential assembly fragmentation. The presence of this contig was confirmed in the mite sample, pineapple O1, using RT-PCR assays (Table 3). Since this contig exhibited low read abundance (Table 2; Figure 2B), and as discussed below because only a partial fragment was recovered, this contig was designated Dolichotetranychus-associated cile-like virus contig (Figure 3), considering its origin from *Dolichotetranychus* mites infesting pineapple. Moreover, two contig sequences (3,511 and 3,585 bp) found in the hibiscus dataset were predicted to code for single ORFs coding for putative proteins of 126 and 125 kDa, respectively. The putative 126 kDa protein was predicted to contain MET and HEL domains, while the putative 125 kDa protein contained MET and RdRp domains (Figure 3). BLASTx searches revealed that the putative 126 and 125 kDa proteins presented ∼30-32% identity to Bemisia tabaci bromo-like virus 3 (QWC36511), and ∼42.8% identity to Erysiphe necator associated bluner-like virus 1 (QKS69535), respectively. Considering the resemblance of these two contigs to RNA 1 and 2 of blunerviruses (Morozov et al., 2020; Figure 3), the contigs were named *Brevipalpus*-associated bluner-like virus RNA 1 and 2. The two bluner-like virus contigs were confirmed in the mite sample hibiscus O1 using RT-PCR assays (Table 3).

#### Contig sequences representing putative new viruses related to negeviruses

Two contig sequences of 9,664 bp and 2,089 bp were *de novo* assembled from the hibiscus and pineapple-coffee mite datasets. Using RT-PCR assays, both contig sequences were confirmed in the mite samples hibiscus O3 and pineapple O1, respectively (Table 3). BLASTx searches showed that both contigs shared 41.9% protein identity with 55% query coverage to Fort Crokett virus (YP_009351834) and 34.3% protein identity with 19% query coverage to Wallerfield virus (AIS40857), respectively. Based on these low sequence identities and their phylogenetic placement (see below), the 9,664 bp and 2,089 bp contigs were considered to represent putative new viruses and were designated Brevipalpus-associated negevirus and Dolichotetranychus-associated negevirus contig, respectively. Similar to the cile-like sequence above, this negevirus-like contig showed low read abundance across the library (Table 2; Figure 2B), consistent with representation from only a small number of mites. Because this sequence represents only a partial genomic fragment, it is referred to here as a putative negevirus-like contig without formal taxonomic assignment. Brevipalpus-associated negevirus was predicted to contain three large ORFs putatively coding for 275, 45, and 23 kDa proteins, respectively (Figure 3). The putative 275 kDa protein was predicted as a replication-associated polyprotein containing multiple protein domains involved in virus replication: MET, methyltransferase FtsJ (FtsJ), HEL, and RdRp. The putative 23 kDa protein was predicted to have the virion membrane protein of plant and insect viruses conserved domain (SP24), while no protein domain was found in the putative 45 kDa protein. In addition, two small overlapping ORFs (<300 bp) were detected within the region encoding the replication-associated polyprotein; both encoded predicted proteins of 4-5 kDa containing at least one transmembrane domain. For Dolichotetranychus-associated negevirus contig, several small ORFs were predicted; however, only a putative protein encoded by an 838 bp ORF presented homology to Wallerfield virus using BLASTx searches. For descriptive purposes, a single continuous ORF was inferred to represent this region, acknowledging possible assembly fragmentation. A chroparavirus MET domain (PFAM 19223) was found within this ORF (Figure 3).

### Phylogenetic relationships of the tenuipalpid mite populations and putative new negeviruses and kitavirids found within the tenuipalpid-associated virome

#### Phylogenetic relationships of the tenuipalpid mite populations inferred using the 28S rRNA and COI genes

Phylogenetic relationships of the tenuipalpid mite populations were inferred using multiple nucleotide sequence alignments of the 28S rRNA and COI genes and the Maximum Likelihood algorithm. Overall, both barcoding markers supported similar phylogenetic relationships among the major *Brevipalpus* lineages. The phylogenies for *B. obovatus* and *B. azores* were identical in both 28S rRNA- and COI-inferred relationships and were placed in single monophyletic lineages specific to each species. Marker-specific differences were observed for *B. yothersi* and *B. papayensis*. Although both species were placed in single monophyletic clades in the 28S rRNA phylogeny, two main clades were observed within the lineages specific for both *Brevipalpus* species (Figures 4 and S1). Notably, the *B. papayensis* clade was paraphyletic in the COI phylogeny, contrasting with the monophyletic *B. papayensis* clade observed in the 28S rRNA-inferred phylogeny. In the COI phylogeny, sequences assigned to *B. azores* formed a distinct clade nested within the broader, paraphyletic *B. papayensis* lineage (Figure 4). Interestingly, although one partial 28S rRNA gene sequence that was amplified from the coffee O1 mite sample presented 98.48% nucleotide identity to *B. papayensis* (MT664798, Table S1), this sequence was placed in a clade that was basal to the lineage containing the *B. papayensis*, *B. feresi, B. phoenicis,* and *B. azores* clades in the 28S rRNA phylogeny (Figure S1). Except for this sequence, all barcoding sequences from *Brevipalpus* mite samples clustered consistently within the *B. yothersi*, *B. papayensis*, *B. azores*, and *B. obovatus* clades in both 28S rRNA and COI phylogenies. Furthermore, although no *Dolichotetranychus* homolog was found in the GenBank database for the 28S rRNA phylogeny, the partial COI sequence from the pineapple O1 mite sample was grouped with another accession corresponding to *Dolichotetranychus* specimens retrieved from pineapple (MH606191). These results corroborate the suggested identities of all the inferred tenuipalpid mites using BLASTn searches (Tables S1 and 2).

**Figure 4.**
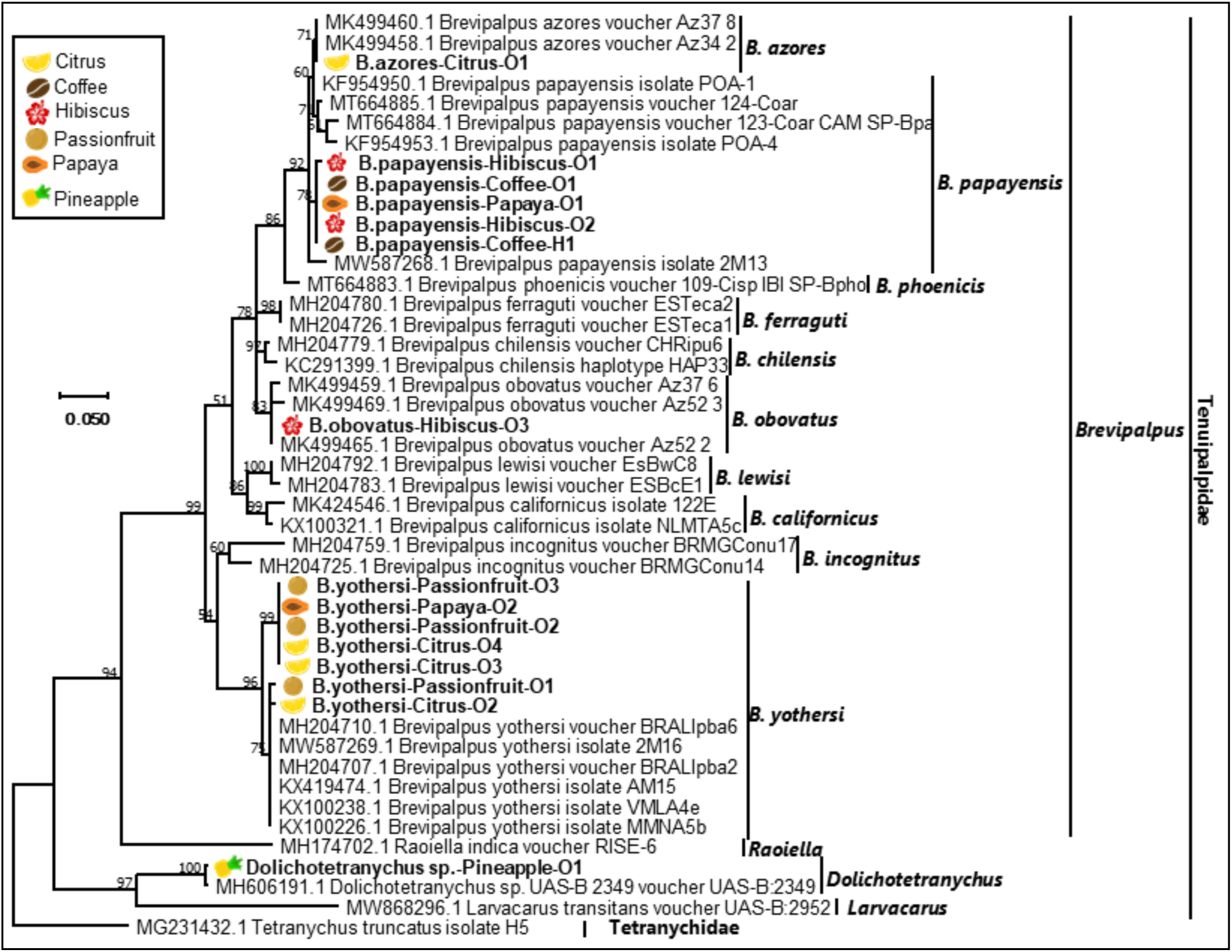
Phylogenetic relationships using partial cytochrome oxidase I (COI) gene sequences from several tenuipalpid mites classified as *Brevipalpus, Raioella, Dolichotetranychus* and *Larvacarus*. Relationships were based on a multiple nucleotide sequence alignment using CLUSTAL and inferred using the Maximum Likelihood algorithm implemented in MEGA 7.0.25. Bootstrap values greater than 50 are shown above the branches after 1,000 repetitions. A homolog partial COI gene sequence from *Tetranychus truncatus* (Tetranychidae) was used as an outgroup. The scale at left indicates the number of substitutions per given branch length. Tenuipalpid mite samples used in this study are bold and placed next to the respective icon of the plant/fruit from which they were collected.

#### Phylogenetic relationships of the putative new negeviruses and kitavirids found within the tenuipalpid mite-associated virome

Phylogenetic relationships of the kitavirids and putative new negeviruses were inferred using multiple protein sequence alignments of the RdRp, HEL, and MET protein domains. Although similar tree topologies and evolutionary relationships were inferred from these three protein domains (Figures 5, S2, and S3), a few notable differences were found. The kitavirids, CiLV-C2H and HGSV-2, were grouped within a clade containing cileviruses, cile-like viruses, and higrevirus, in the three conserved protein domains. They were grouped with the previously characterized isolates of the same virus species (CiLV-C2, ATW76030; HGSV-2, AER13445). Similarly, the putative new negevirus found in the mite sample hibiscus O3: Brevipalpus-associated negevirus, clustered with Fort Crockett virus (YP009351834) in the three phylogenies. The clade containing the two previously mentioned negeviruses shared a common evolutionary origin with the recently reported group ‘aphiglyvirus’ (Kondo et al., 2020) within the unofficial negevirus taxon (Figures 5, S2, and S3).

**Figure 5.**
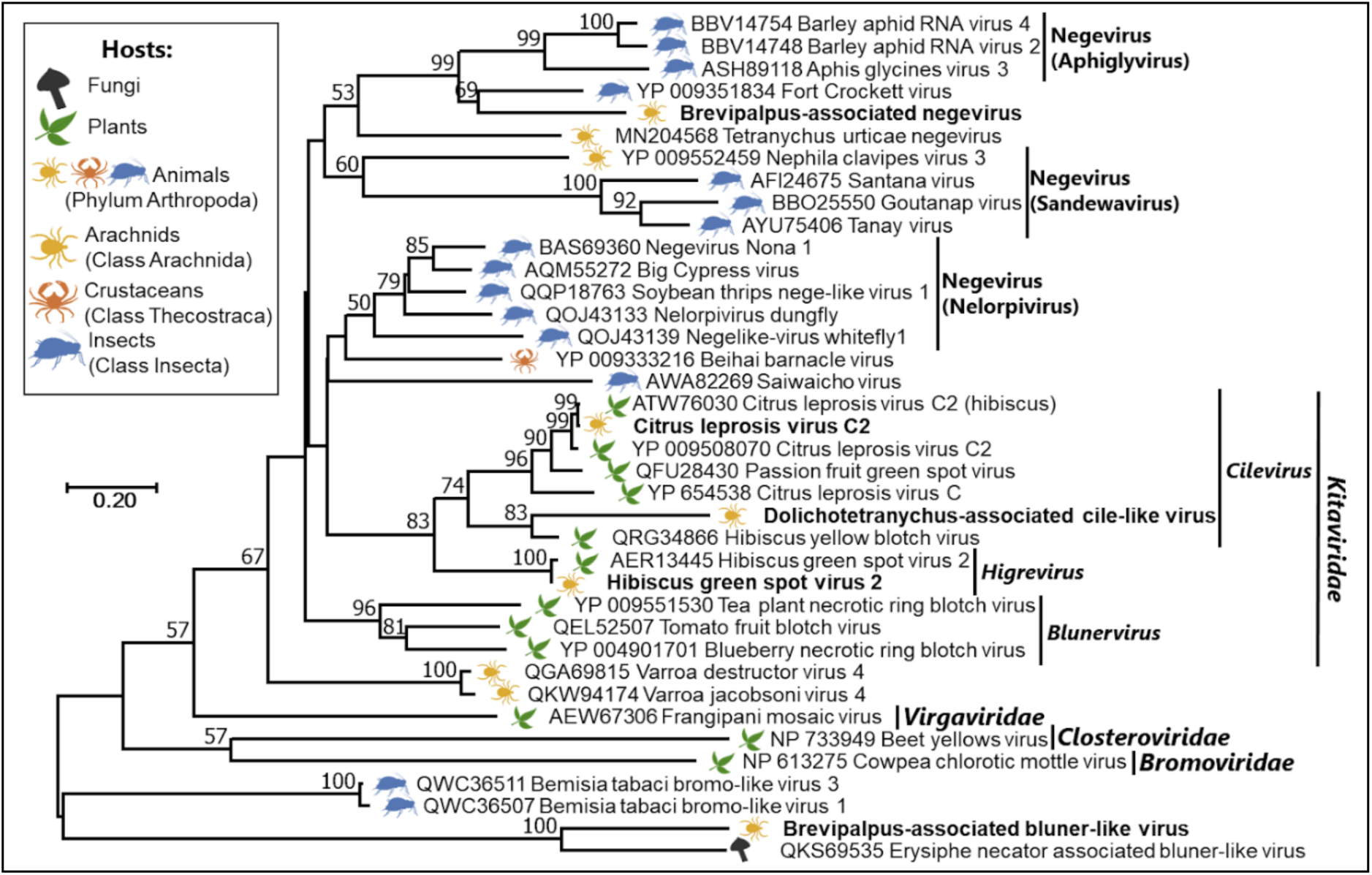
Phylogenetic relationships inferred using the RNA-dependent RNA polymerase (RdRp) conserved domain found in the contig sequences of the putative new negeviruses and kitavirids in tenuipalpid mite samples. The specific mite sample from which each putative new virus was detected is detailed in Table 3. Brevipalpus-associated bluner-like virus corresponds to the RdRp domain present in the Brevipalpus-associated bluner-like virus contig 2 (RNA 2) (Figure 3). Dolichotetranychus-associated cile-like virus clustered within the polyphyletic *Kitaviridae* clade in the RdRp phylogeny. No clear evolutionary relationships were inferred for Brevipalpus-associated bluner-like virus.

### Diversity measurement of flat mites virome

Diversity of flat mites virome within and across different samples were examined. Alpha diversity matrixes showed range of values across five different pools (Table S3). Viral richness (number of taxonomic groups detected per pool) ranged from 2 to 6. The highest richness was observed in mites from hibiscus, while mites from passionfruits contained the fewest taxa. Shannon diversity indices, based on normalized abundance, varied between 0.19 in Citrus to 1.41 in hibiscus, showing that some pools, such as citrus and papaya, were dominated by a few viral families, while others, such as hibiscus and pineapple, exhibited a more even community structure. Community similarity among pools was evaluated using both taxonomic count data and normalized abundance data, based on Jaccard or Bray–Curtis dissimilarities (β-diversity; “similarity” reported as 1 – dissimilarity values) (Figure S4). Ordination with NMDS produced a stress value near zero, which reflects the small number of pools and limits the interpretability of the ordination (Figure 6). Across both datasets, β-diversity measurements and NMDS plots indicated that viral communities were distinct among pools. Jaccard similarity ranged from 0–0.67, and Bray–Curtis similarity ranged from 0–0.36. In Jaccard comparisons, citrus and hibiscus pools were the most similar, sharing four taxonomic groups: *Narnaviridae*, *Ourmiaviridae*, *Picornavirales*, and *Tombusviridae*. In Bray–Curtis comparisons, hibiscus and papaya pools showed the highest similarity, driven by the presence of *Picornavirales* and *Kitaviridae* in both pools. Overall, each pool represents unique virome structure, which highlights the diversity of flat mites associated virome.

**Figure 6.**
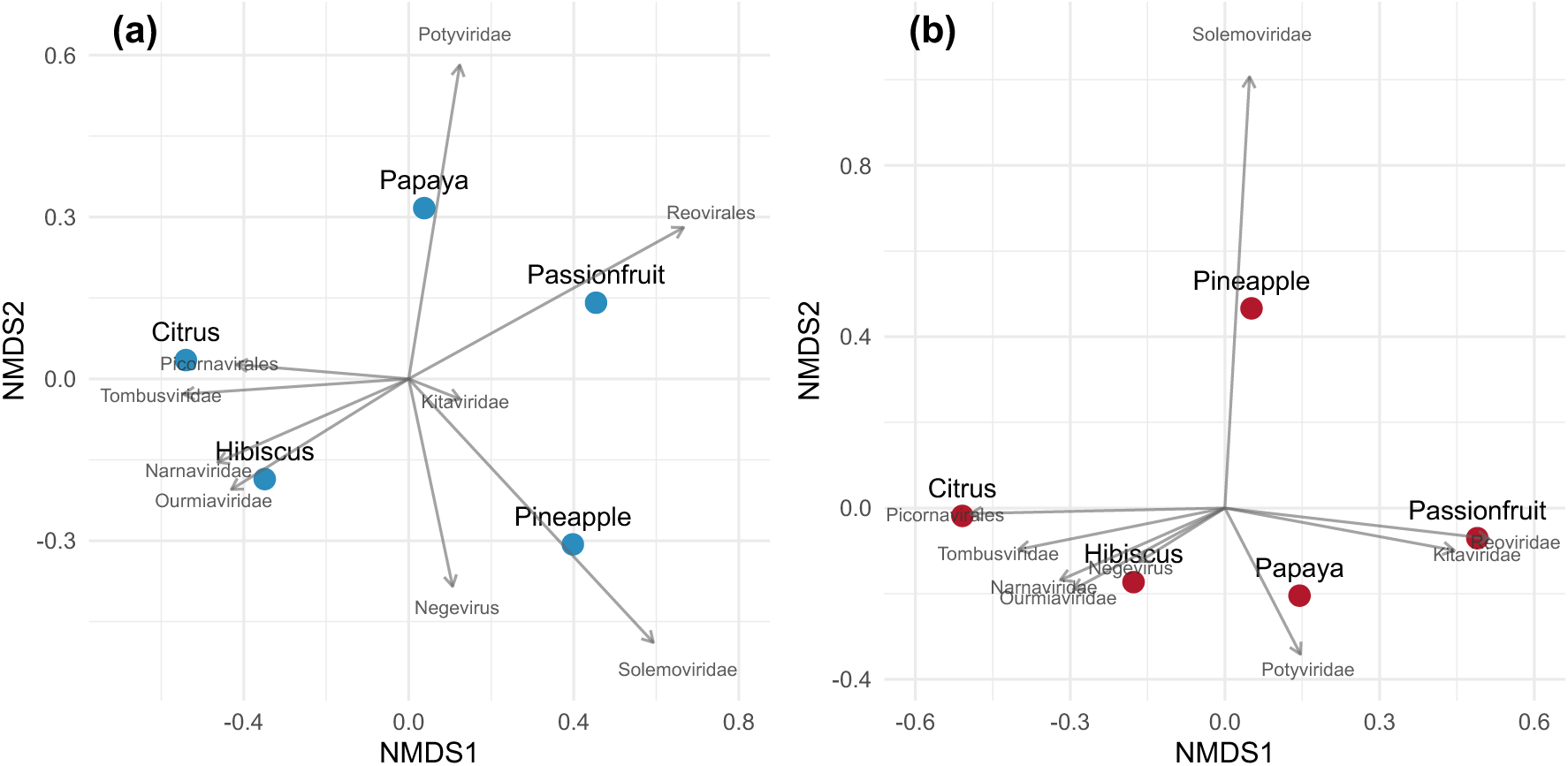
Non-metric multidimensional scaling (NMDS) ordination of virome composition based on (a) the number of viral species detected per taxonomic group (count dataset) and (b) the total read abundance per taxonomic group (abundance dataset). Each colored point represents an individual virome pool. Vectors indicate viral taxonomic groups and their directional association with the ordination axes; arrow length reflects the strength of correlation between each group and community structure.

## DISCUSSION

This study describes the characterization of kitavirus-related viruses within the diverse virome found in flat mite specimens collected from several hosts on the Oahu and Hawaii Islands.

Using a CTAB-based TNA extraction protocol (Li et al., 2008), followed by column purification, was critical for recovering high-quality TNA from as little as 10 mites. Although ribodepleted total RNA is currently considered one of the best templates, if not the best, for transcriptomics and virome studies in plant and arthropod samples (Villamor et al., 2019; Fauver et al., 2019), TNA extracts proved helpful in studying the virome present in the mite samples in this study. Despite the inclusion of a ribodepletion step during library construction, abundant rRNA sequences were still recovered (data not shown).

Accordingly, standard rRNA depletion protocols may perform poorly in non-model arthropods. In future studies, ribosomal RNA depletion from RNA or TNA samples from non-model animal hosts may succeed better using class- or family-specific probes and a reverse-transcription/RNAse-H based protocol (Fauver et al., 2019). Previous HTS studies using ribodepleted total RNA as a template have established a suggested minimum sequencing depth of 0.5 – 10 M of reads for virus diagnostics and discovery (Massart et al., 2018; Bester et al., 2021; Huang et al., 2019; Malapi-Wight et al., 2021). Therefore, the high sequencing depth used in this study, ranging from ∼65-91 M reads (Table 1), likely compensated for inefficient ribodepletion. Regardless, the library construction method used in this study, the number of raw reads obtained per dataset, and bioinformatic analyses allowed for characterization of the flat mite virome, including the identified BTV-C and BTV-C-like viruses.

After removing as many reads with tenuipalpid origin as possible, a variable number of non-host reads (∼21 – 62 M of reads) were *de novo* assembled (Table 1). Although less than 0.5% of the non-host reads were of viral origin, assembly of these reads yielded contigs representing, in some cases, the near-complete genome of CiLV-C2H, HGSV-2, and several putative new virus species. These sequences are interpreted here as mite-associated viral sequences detected in mite-derived nucleic acid extracts, classified within the *Picornavirales* order, *Narnaviridae*, *Tombusviridae*, *Kitaviridae*, *Solemoviridae,* and *Potyviridae* families, and the unofficial Negevirus taxon (Figures 2-3 and data not shown). Picornavirids, tombusvirids, narnavirids, reovirids, solemovirids, and negeviruses have been similarly characterized from other arthropods and insects (Debat, 2017; Guo et al., 2021b; Huang et al., 2021). The presence of the putative new viruses was validated in at least one mite sample using RT-PCR assays, except for one narnavirus and two picornaviruses, which could not be detected in the hibiscus mite samples (Table 3). It is possible that the concentration of those putative new viruses was below the sensitivity achieved using RT-PCR assays. Alternatively, these contigs may represent contamination introduced during library preparation or *in silico* demultiplexing, which are recognized sources of HTS artifacts (Lee et al., 2016; Lusk, 2014).Although BTV-N-related viruses, namely rhabdovirids, have been characterized from other arthropods, including ticks and mites (Sameroff et al., 2019; Guo et al., 2021b), no rhabdovirid or negative-sense RNA virus was found in the datasets generated here. The predominance of RNA viruses is consistent with other arthropod virome studies, in which positive-sense RNA viruses frequently dominate (Shi et al., 2016). Importantly, ribodepleted TNA/RNA libraries are not inherently restricted to RNA virus detection, as DNA viruses can be recovered when transcriptionally active, as demonstrated in both plant and arthropod systems (Soltani et al., 2021; Larrea-Sarmiento et al., 2024). Therefore, the absence of DNA viruses and negative-sense RNA viruses, including the BTV orchid fleck virus (OFV), likely reflects low local prevalence and sampling context rather than a limitation of the extraction or sequencing strategy.

We also note that the choice of bioinformatic pipeline influences virome interpretation. We implemented a BLASTx-based identification pipeline and validated viral contigs by identifying viral hallmark genes and conserved domains, assessing ORF organization, and confirming read-mapping support. In contrast, an alternative pipeline using VirSorter2 did not identify more than half of these viral sequences, although it detected two additional ourmiavirus-like contigs not identified by the BLASTx-based approach.

Picornavirid sequences were found in three of the four *Brevipalpus*-derived viral datasets from citrus, hibiscus, and papaya hosts (Figure 2), and they represented the most abundant viral group in the citrus dataset. Similar dominance of picornavirids has been reported in the virome of nephilid spiders (Debat, 2017). Because picornavirids in arthropods can cause outcomes ranging from asymptomatic infection to paralysis and mortality (van Oers, 2010; Valles et al., 2017), further molecular and biological characterization of these viruses in *Brevipalpus* mites is warranted, including evaluation of their potential for biological control.

Near-complete genomes of CiLV-C2H were retrieved from the passionfruit and papaya mite datasets (Figure 3) and confirmed by RT-PCR assays in individual mite samples (Table 3). Although viruliferous *B. yothersi* mites carrying CiLV-C2H were collected from papaya leaves, no obvious viral-like symptoms were observed. This suggests the incapability of CiLV-C2H to infect papaya, or obvious symptoms may be only visible once leaves senesce (Ramos-Gonzalez et al., 2020; Melzer et al., 2012; Melzer et al., 2013; Roy et al., 2015; Olmedo-Velarde et al., 2021). Barcoding these mite samples using the 28S rRNA and COI genes and phylogenetic analyses suggested their identity as *B. yothersi* (Tables 1 and S1, and Figures 4 and S1). While *B. yothersi* has been demonstrated to transmit CiLV-C2 (Roy et al., 2013), and related *Brevipalpus* species are competent vectors of multiple kitavirids, the detection of CiLV-C2H RNA in *B. yothersi* in this study indicates virus acquisition, rather than confirmed transmission, which will require adequate transmission assays.

A near-complete genome of HGSV-2 was retrieved from the hibiscus mite dataset (Figure 3). The genomic organization was identical to that obtained from hibiscus and citrus HGSV-2 isolates from Oahu and Maui Islands, respectively (Olmedo-Velarde et al., 2024). The presence of HGSV-2 was confirmed in the hibiscus O1 sample using RT-PCR assays (Table 3). Barcoding this mite sample using the 28S rRNA and COI genes and phylogenetic analyses suggested its identity as *B. papayensis* mites (Table 1, Figures 4 and S1). Transmission assays of HGSV-2 using mites from a *B. azores* colony demonstrated the ability of this *Brevipalpus* species to transmit HGSV-2, although with low efficiency (Olmedo-Velarde et al., 2024). More recently, experimental transmission of HGSV-2 by both *B. yothersi* and *B. papayensis* was demonstrated under controlled conditions, confirming vector competence of these species (Pereira et al., 2025). Considering the close evolutionary relationship between *B. papayensis* and *B. azores*, which are placed in the same phylogenetic clade of 28S rRNA and COI phylogenies (Figures 4 and S1), the detection of HGSV-2 RNA in field-collected *B. papayensis* mites in this study is consistent with these experimental transmission studies. However, experimental confirmation of transmission using the HGSV-2 isolate from Hawaii will be required.

Although overall topologies inferred from 28S rRNA and COI were largely congruent, minor inconsistencies between markers were observed, particularly in samples where more than one *Brevipalpus* species was detected. Such discrepancies may reflect differences in phylogenetic resolution between markers or primer-dependent amplification bias favoring specific species haplotypes, rather than biological incongruence. Improving taxonomic resolution for *Brevipalpus* and *Dolichotetranychus* will likely require integrative approaches combining multiple genetic markers with detailed morphological validation.

Interestingly, several contig sequences corresponding to the genome of PRSV were also found in the papaya mite dataset, and their presence was confirmed by RT-PCR (Table 3). Because tenuipalpid mites are phytophagous, the detection of plant virus–related sequences may reflect just ingestion from infected tissue rather than active replication or vector competence within the mite.

Near-complete genomes of a putative new negevirus, Brevipalpus-associated negevirus, and a putative new bluner-like virus, Brevipalpus-associated bluner-like virus, were retrieved from the hibiscus mite dataset. Partial genomic sequences of a putative new negevirus, Dolichotetranychus-associated negevirus contig, and a putative new cile-like virus, Dolichotetranychus-associated cile-like virus contig, were retrieved from the pineapple-coffee mite dataset (Figure 3), and their presence was confirmed by RT-PCR. Brevipalpus-associated negevirus (∼9.6 Kb), detected in the sample hibiscus O3 (Table 3), presented the genomic organization and conserved domains typical of negeviruses (Vasilakis et al., 2013; Nunes et al, 2017). Low protein identity and divergence from other negeviruses, including Fort Crocket virus and aphiglyviruses in RdRp, HEL, and MET phylogenies (Figures 5, S2, and S3), suggest Brevipalpus-associated negevirus represents a putative new negevirus most closely related to aphiglyviruses. Negevirus genomes typically encode three ORFs: a replication-associated polyprotein, a putative glycoprotein, and a membrane-bound protein containing the SP24 domain, a conserved small transmembrane protein characteristic of negeviruses and kitavirids. Furthermore, the putative glycoprotein contains transmembrane domains and signal peptides similar to those of the cilevirus p61 protein (Kuchibhatla et al., 2014). Although negeviruses and kitavirids have been proposed to share a common ancestor that may have colonized flat mites (Freitas-Astúa et al., 2018; Ramos-González et al., 2020; Quito-Avila et al., 2020), no negevirus had previously been reported from *Brevipalpus* mites. Thus, identification of Brevipalpus-associated negevirus is consistent with proposed evolutionary scenarios between negeviruses and kitavirids. However, given the limited taxon sampling and the absence of comprehensive phylogenomic analyses, these findings should be interpreted as supportive rather than definitive. Kitavirid ancestors may have originated from arthropod-infecting nege-like viruses through genome segmentation, recombination, and acquisition of genes for plant cell movement and viral suppression of RNA silencing. These gene acquisitions likely occurred from plant RNA viruses within the order *Martellivirales* (Ramos-González et al., 2021; Dolja et al., 2020), although additional comparative genomic data are required to robustly evaluate this model.

Brevipalpus-associated bluner-like virus (RNA 1, 3,511 bp; RNA 2, 3,585 bp) exhibited a genomic organization resembling that of blunerviruses, with one notable difference. In blunerviruses, RNA 1 encodes a polyprotein containing MET and HEL domains, whereas RNA 2 encodes a polyprotein containing HEL and RdRp domains (Figure 3) (Quito-Avila et al., 2020). In constrast, the RNA 2-encoded polyprotein of Brevipalpus-associated bluner-like virus contained a MET rather than a HEL domain (Figure 3). Low protein identity and divergence from viruses classified within the *Martellivirales* order in RdRp, HEL, and MET phylogenies (Figures 5, S2, and S3), indicate that virus is highly divergent that its precise taxonomic placement within *Martellivirales* remains unresolved. Given the recent experimental transmission of blunerviruses by eriophyid mites (Tassi et al., 2025), virome characterization of eriophyid mites will be important for testing evolutionary scenarios linking arthropod-and plant-associated viruses. Koonin et al. (2021) proposed a conceptual framework describing the virosphere as a dynamic continuum comprising the orthovirosphere, viruses with both replication and structural protein genes, and the perivirosphere that has virus-like agents encoding replication-associated proteins but lacking recognizable structural protein genes. Based on the absence of an identifiable capsid protein gene, Brevipalpus-associated bluner-like virus may tentatively be placed within the perivirosphere of *Martellivirales*. Nevertheless, the presence of an additional genomic segment encoding a highly divergent capsid protein cannot be excluded.

Small ORFs encoding orphan transmembrane proteins were abundant in the genomes of Brevipalpus-associated negevirus and Brevipalpus-associated bluner-like virus (Figure 3), consistent with observations from other negeviruses and kitavirids (Kuchibhatla et al., 2014; Ramos-González et al., 2021). These proteins may contribute to regulation of host cellular processes and increased virus fitness (Dimaio, 2014).

Finally, partial sequences of Dolichotetranychus-associated negevirus contig and Dolichotetranychus-associated cile-like virus contig, both detected in the sample pineapple O1 (Table 3), contained MET and RdRp conserved domains, respectively (Figure 3). Although low protein identity and phylogenetic placement suggest these sequences represent putative new viruses, further work is required to fully characterize them. The low number of reads mapping to these sequences (Figure 2), suggest that a small subset of *Dolichotetranychus* mites in the pool carried these viruses, limiting recovery of complete genomes despite deep sequencing (>65 M reads per pool). Characterization of Dolichotetranychus-associated cile-like virus and evaluation of virus colonization in pineapple tissue is of great interest for the pineapple industry, considering the cile-like virus nature and the importance of cileviruses in causing plant diseases of economic significance.

In conclusion, this study establishes a baseline framework for flat mite virome surveillance, enabling detection of BTVs that may emerge as agriculturally important pathogens and providing insight into BTV evolution. Given the limited number of samples and hosts examined here, and the broad host range of flat mites, substantial opportunities remain to characterize additional BTVs and BTV-related viruses in *Brevipalpus* and other Tenuipalpidae populations worldwide.

## FUNDING

This project was funded, in part, by USDA-NIFA Hatch Project HAW09050-H awarded to MM and managed by the College of Tropical Agriculture and Human Resources at the University of Hawaii at Manoa.

## Supplementary Material

**Table S1.**
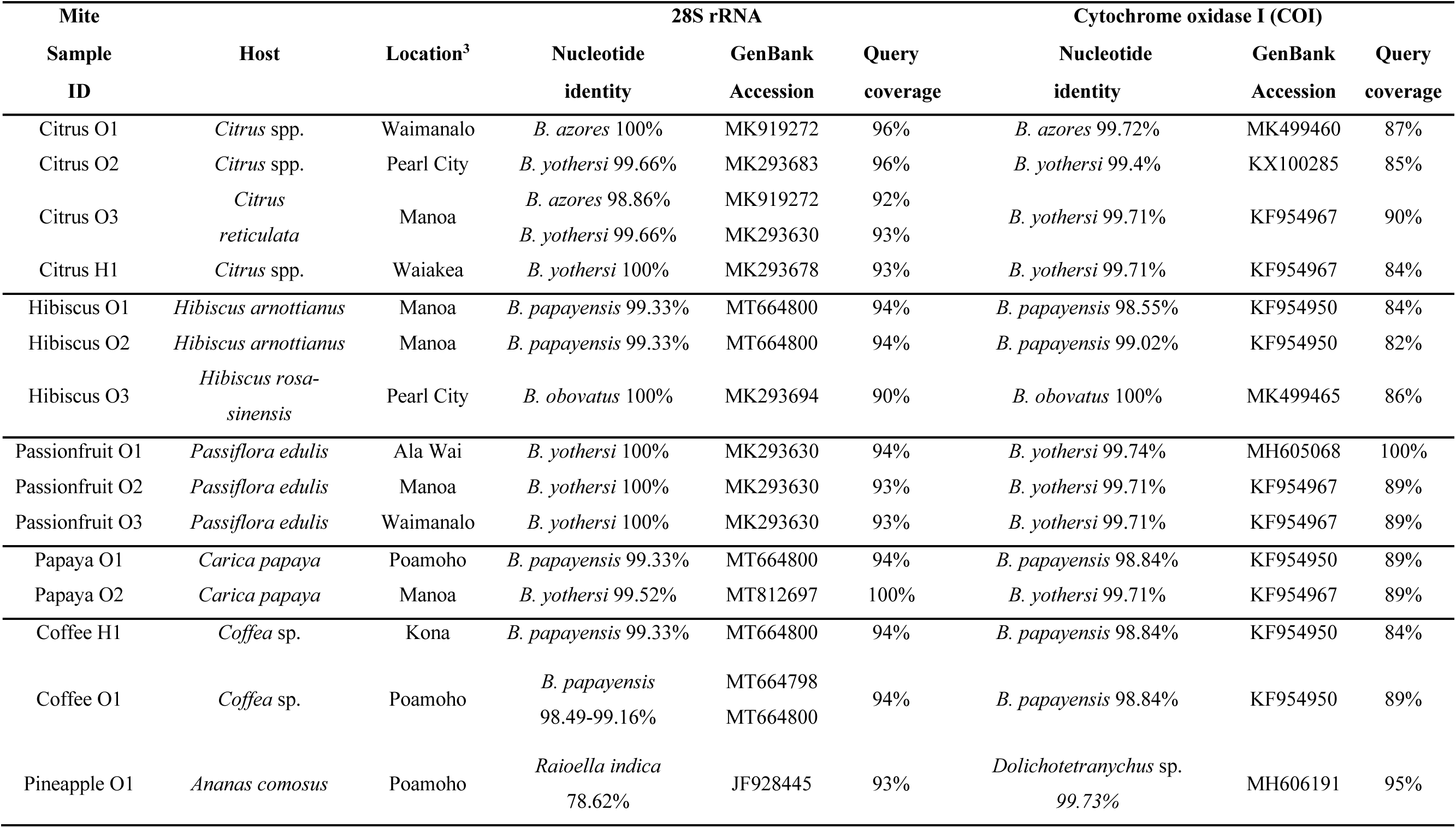
BLASTn searches results of the partial 28S rRNA and COI sequences amplified from the mite samples used in this study.

**Table S2.**
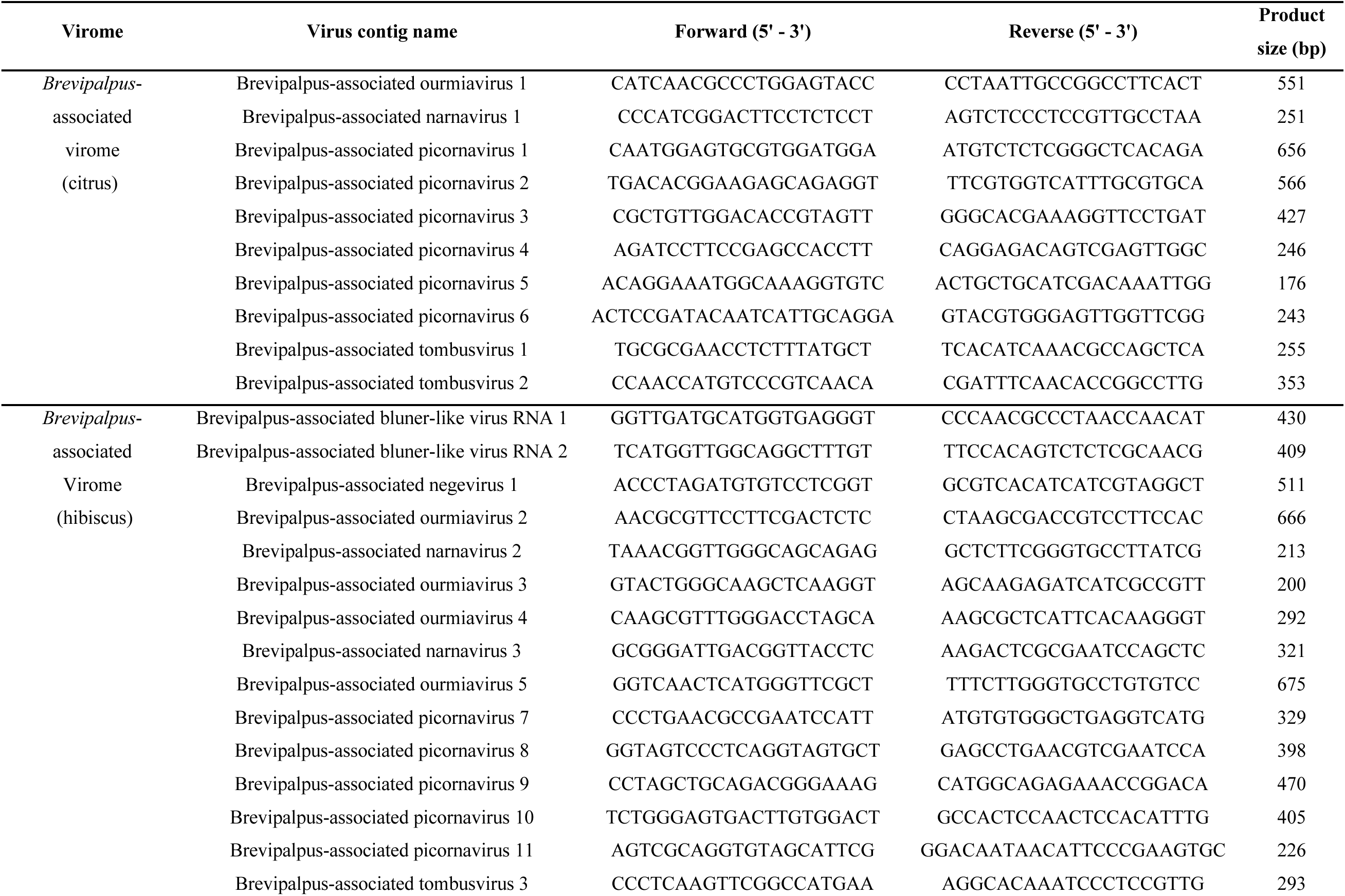
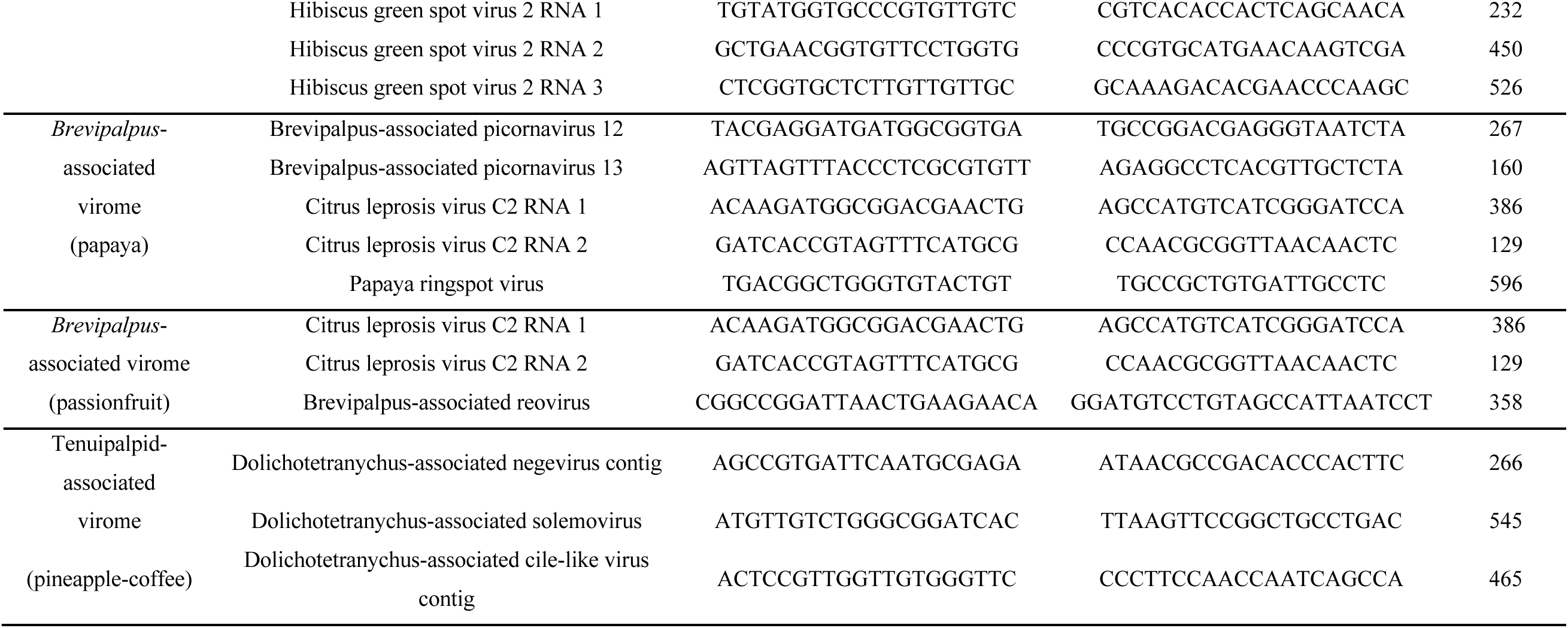
Primer sequences used in RT-PCR assays for virus presence confirmation from the different mite samples used in this study.

**Table S3.**
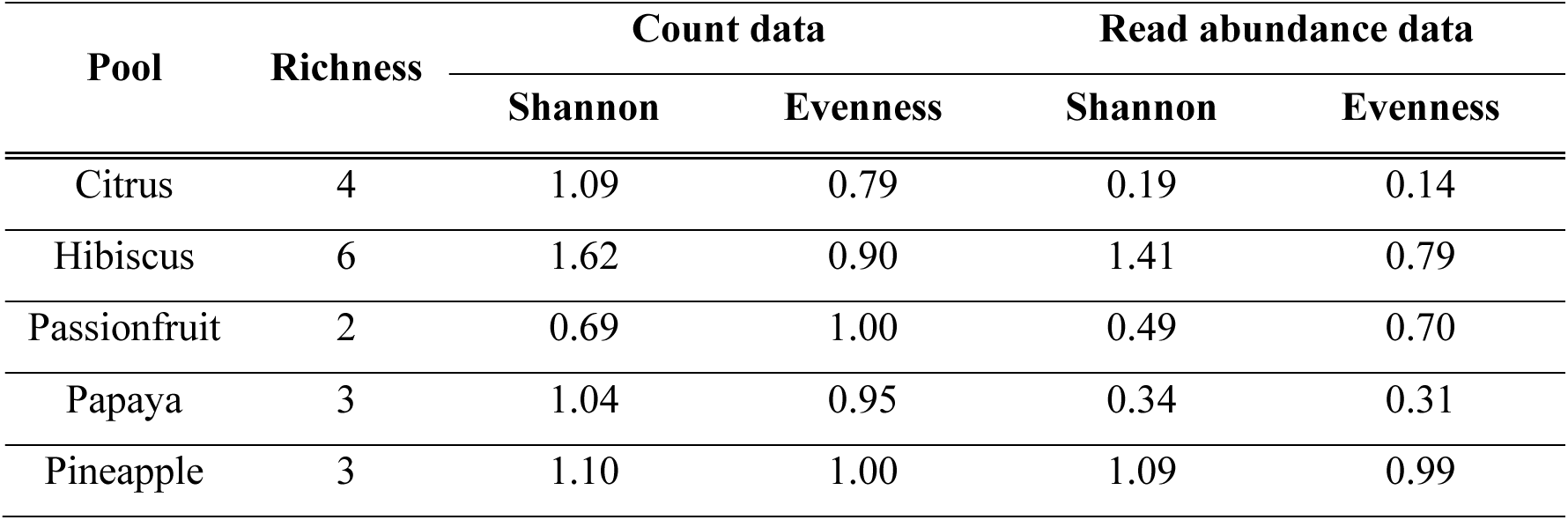
Alpha-diversity metrics of each virome pool. Richness (number of viral taxonomic groups), Shannon diversity, and Pielou’s evenness were calculated separately for the count and read-abundance datasets, each aggregated by assigned viral taxonomic group.

**Figure S1.**
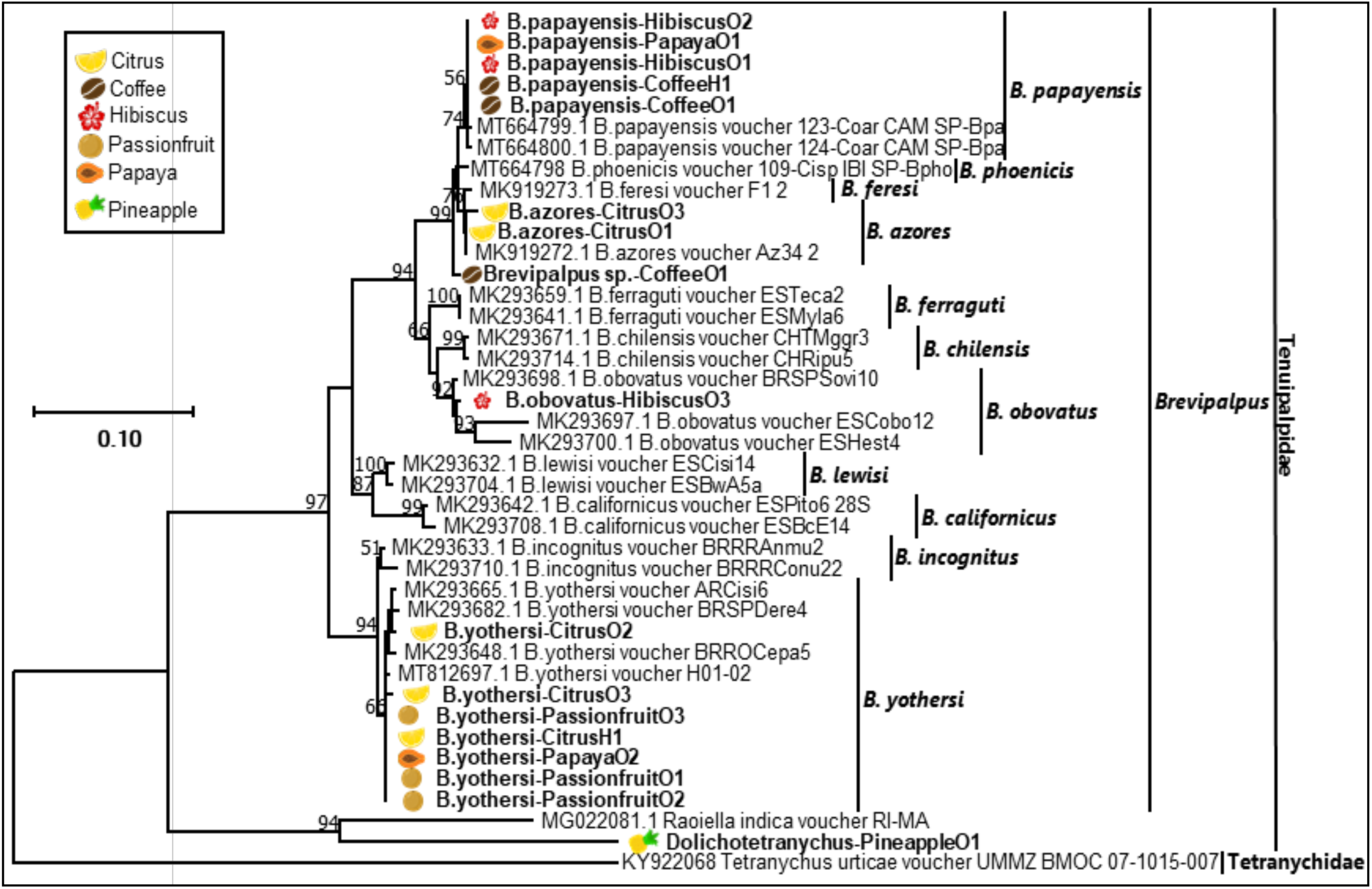
Phylogenetic relationships using partial 28S rRNA gene sequences from several tenuipalpid mites classified within *Brevipalpus*, *Raioella* and *Dolichotetranychus*. Relationships were based on a multiple nucleotide sequence alignment using CLUSTAL and inferred using the Maximum Likelihood algorithm implemented in MEGA 7.0.25. Bootstrap values greater than 50 are shown above the branches after 1,000 repetitions. No *Dolichotetranychus* 28S rRNA sequence was found in the GenBank database. A homolog partial 28S rRNA gene sequence from *Tetranychus urticae* (Tetranychidae) was used as an outgroup. The scale at the left indicates the number of substitutions per given branch length. Tenuipalpid mite samples from in this study are bold and placed next to the respective icon of the plant/fruit from which they were collected.

**Figure S2.**
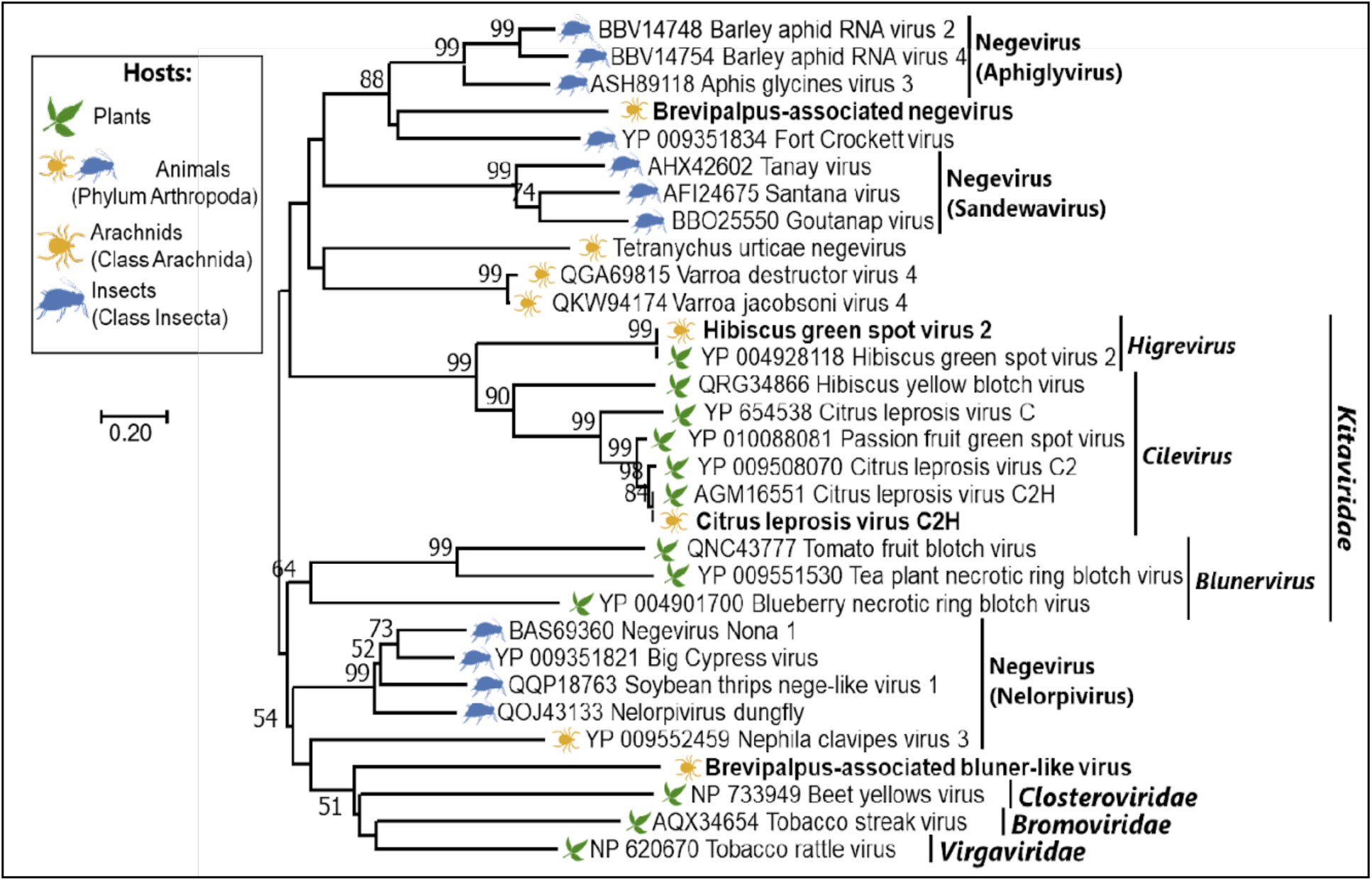
Phylogenetic relationships inferred using helicase (HEL) conserved domain found in the contig sequences of the putative new negeviruses and kitavirids in tenuipalpid mite samples. The specific mite sample that each putative new virus was detected from is detailed in Table 3. Brevipalpus-associated bluner-like virus corresponds to the RdRp domain present in the Brevipalpus-associated bluner-like virus contig 1 (RNA 1) (Figure 3).

**Figure S3.**
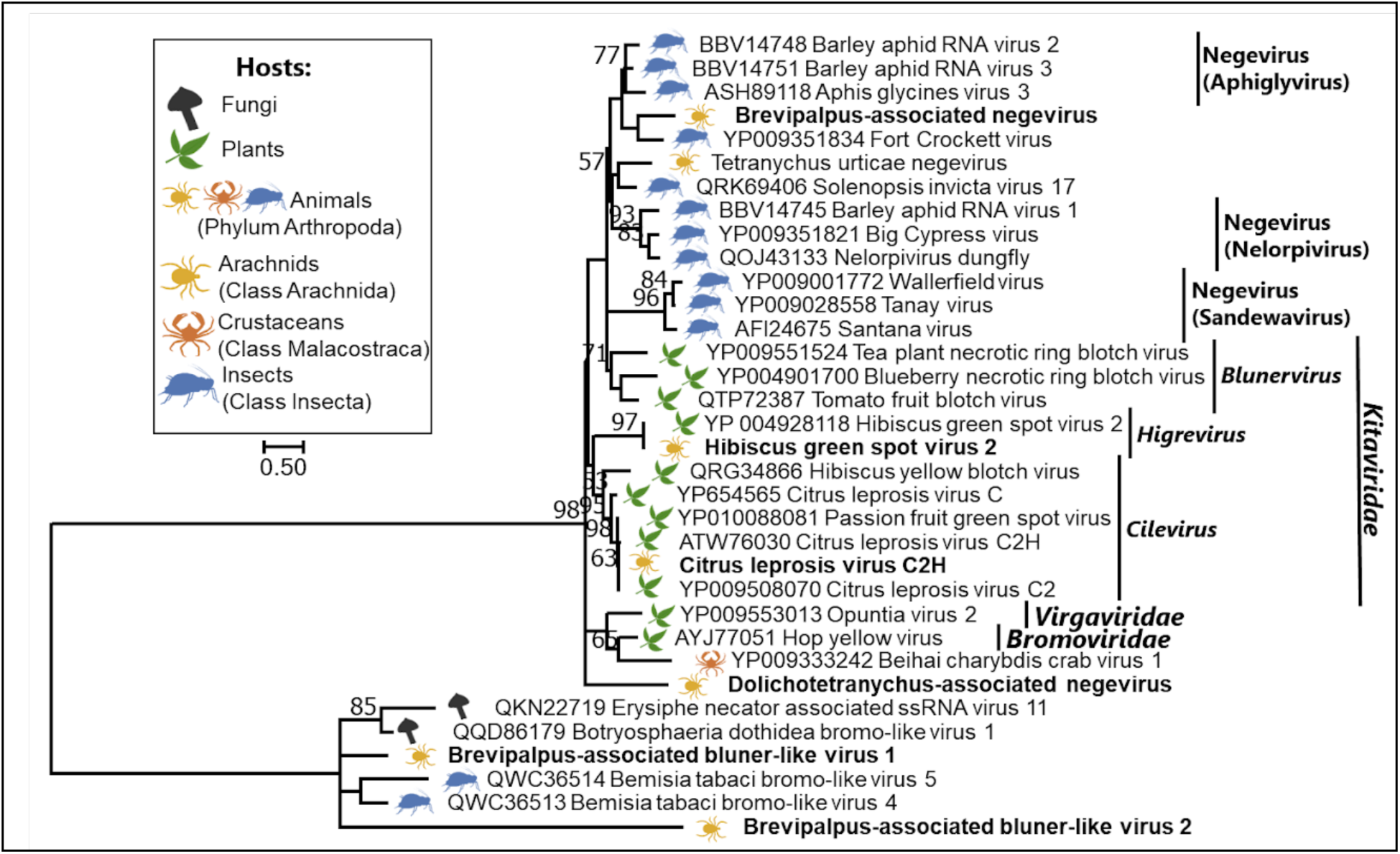
Phylogenetic relationships inferred using the methyltransferase (MET) conserved domain found in the contig sequences of the putative new negeviruses and kitavirids in tenuipalpid mite samples. The specific mite sample that each putative new virus was detected from is detailed in Table 3. Brevipalpus-associated bluner-like virus 1 and 2 correspond to the MET domains present in the Brevipalpus-associated bluner-like virus contig 1 (RNA 1) and 2 (RNA 2), respectively (Figure 3).

**Figure S4.**
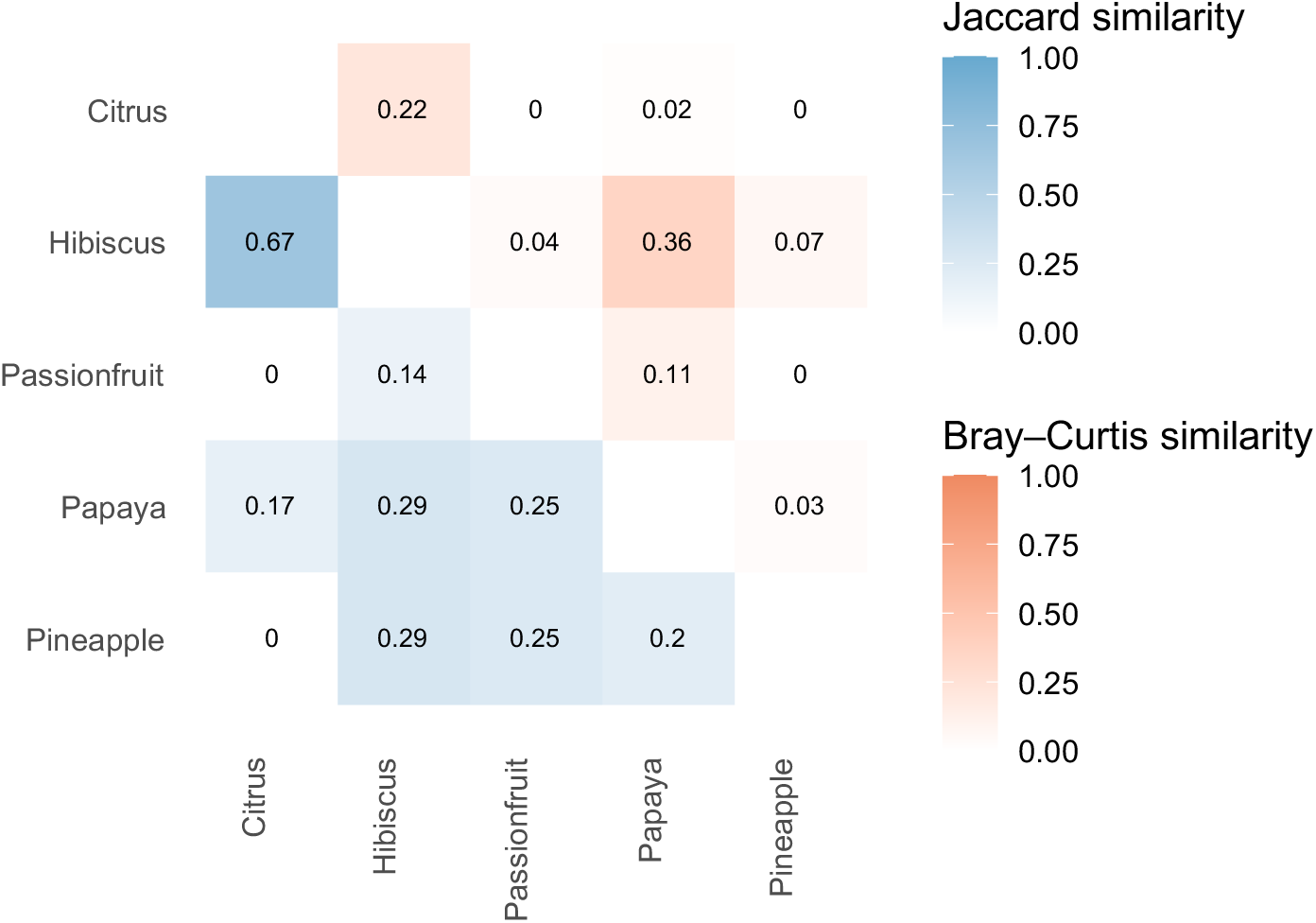
Heatmap showing pairwise β-diversity among virome pools based on Jaccard (blue) and Bray–Curtis (red) similarity metrics. The lower triangle represents Jaccard similarity calculated from the count dataset, and the upper triangle represents Bray–Curtis similarity calculated from the read-abundance dataset. Similarity values were derived as 1 - dissimilarity, where 0 = completely dissimilar and 1 = identical.

## Notes

### Competing Interest Statement

The authors have declared no competing interest.

## LITERATURE CITED

Altschul, S.F., Madden, T.L., Schäffer, A.A., Zhang, J., Zhang, Z., Miller, W., and Lipman, D.J. 1997. Gapped BLAST and PSI-BLAST: a new generation of protein database search programs. Nucleic Acids Res. 25:3389–3402.

Arif, M. & Ochoa-Corona, F. (2013). Comparative assessment of 5’ A/T-rich overhang sequences with optimal and sub-optimal primers to increase PCR yields and sensitivity. Mol. Biotechnol. 55, 17–26. doi: 10.1007/s12033-012-9617-5

Bankevich, A., Nurk, S., Antipov, D., Gurevich, A. A., Dvorkin, M., Kulikov, A. S., Lesin, V. M., Nikolenko, S. I., Pham, S., Prjibelski, A. D., Pyshkin, A. V., Sirotkin, A. V., Vyahhi, N., Tesler, G., Alekseyev, M. A., & Pevzner, P. A. (2012). SPAdes: a new genome assembly algorithm and its applications to single-cell sequencing. Journal of computational biology: a journal of computational molecular cell biology, 19(5), 455–477. 10.1089/cmb.2012.0021

Beard, J. J., Ochoa, R., Bauchan, G. R., Trice, M. D., Redford, A. J., Walters, T. W., et al. (2013). Flat Mites of the World, 2 Edn. Fort Collins, CO: CPHST.

Beard, J. J., Ochoa, R., Braswell, W. E., and Bauchan, G. R. (2015). Brevipalpus phoenicis (Geijskes) species complex (Acari: Tenuipalpidae)-a closer look. Zootaxa 3944:1–67.

Bester, R., Cook, G., Breytenbach, J.H.J., Steyn, C., De Bruyn, R., Maree, H.J. (2021). Towards the validation of high-throughput sequencing (HTS) for routine plant virus diagnostics: measurement of variation linked to HTS detection of citrus viruses and viroids. Virol J 18, 61. 10.1186/s12985-021-01523-1

Bolger, A. M., Lohse, M., & Usadel, B. (2014). Trimmomatic: a flexible trimmer for Illumina sequence data. Bioinformatics (Oxford, England), 30(15), 2114–2120. 10.1093/bioinformatics/btu170

Chabi-Jesus, C., Ramos-González, P. L., Tassi, A. D., Rossetto Pereira, L., Bastianel, M., Lau, D., Canale, M. C., Harakava, R., Novelli, V. M., Kitajima, E. W., & Freitas-Astúa, J. (2023). Citrus Bright Spot Virus: A New Dichorhavirus, Transmitted by Brevipalpus azores, Causing Citrus Leprosis Disease in Brazil. Plants, 12(6), 1371. 10.3390/plants12061371

Debat, H.J. (2017). An RNA Virome Associated to the Golden Orb-Weaver Spider Nephila clavipes. Front. Microbiol. 8:2097. doi: 10.3389/fmicb.2017.02097

Dey, K.K., Sugikawa J., Kerr, C., Melzer, M.J. (2019) Air potato (Dioscorea bulbifera) plants displaying virus-like symptoms are co-infected with a novel potyvirus and a novel ampelovirus. Virus Genes 55(1):117–121. doi: 10.1007/s11262-018-1616-6.

Dimaio, D. (2014). Viral miniproteins. Annu. Rev. Microbiol. 68, 21–43. doi: 10.1146/annurev-micro-091313-103727

Dolja, V.V., Krupovic, M., Koonin, E.V. (2020). Deep Roots and Splendid Boughs of the Global Plant Virome. Annu. Rev. Phytopathol. 58(1). doi:10.1146/annurev-phyto-030320-041346

Fauver, J. R., Akter, S., Morales, A., Black, W. C., 4th, Rodriguez, A. D., Stenglein, M. D., Ebel, G. D., & Weger-Lucarelli, J. (2019). A reverse-transcription/RNase H based protocol for depletion of mosquito ribosomal RNA facilitates viral intrahost evolution analysis, transcriptomics and pathogen discovery. Virology 528, 181–197. 10.1016/j.virol.2018.12.020

Freitas-Astúa, J., Ramos-González, P.L., Arena, G.D., Tassi, A.D., & Kitajima, E.W. (2018). *Brevipalpus*-transmitted viruses: parallelism beyond a common vector or convergent evolution of distantly related pathogens? Curr Opin Virol 33: 66–73. 10.1016/j.coviro.2018.07.010

Gerson, U. (2008). The Tenuipalpidae: an under-explored family of plant-feeding mites. Systematic and Applied Acarology, 13(2), 83. doi:10.11158/saa.13.2.1

Guo, J., Bolduc, B., Zayed, A. A., Varsani, A., Dominguez-Huerta, G., Delmont, T. O., Pratama, A. A., Gazitúa, M. C., Vik, D., Sullivan, M. B., Roux, S. (2021a). VirSorter2: a multi-classifier, expert-guided approach to detect diverse DNA and RNA viruses. Microbiome. 9(1):37. doi: 10.1186/s40168-020-00990-y.

Guo, L., Lu, X., Liu, X., Li, P., Wu, J., Xing, F., Peng, H., Xiao, X., Shi, M., Liu, Z., Li, X.-D., Guo, D. (2021b). Metatranscriptomic analysis reveals the virome and viral genomic evolution of medically important mites. J Virol. 95(7):e01686–20. doi: 10.1128/JVI.01686-20.

Huang, B., Jennison, A., Whiley, D. McMahon, M., Hewitson, G., Graham, R., De Jong, A., Warrilow, D. (2019). Illumina sequencing of clinical samples for virus detection in a public health laboratory. Sci Rep 9, 5409. 10.1038/s41598-019-41830-w

Huang, X. & Miller, W. (1991). A time-efficient, linear-space local similarity algorithm. Adv. Appl. Math. 12, 337–357.

Huang, H.J.., Ye, Z.X.., Wang, X. Yan, X.-T., Zhang, Y., He, Y.-J., Qi, Y.-H., Zhang, X.-D., … Li, J.-M. (2021). Diversity and infectivity of the RNA virome among different cryptic species of an agriculturally important insect vector: whitefly Bemisia tabaci. npj Biofilms Microbiomes 7, 43 10.1038/s41522-021-00216-5

Kallies, R., Kopp, A., Zirkel, F., Estrada, A., Gillespie, T. R., Drosten, C., Junglen, S. (2014). Genetic characterization of goutanap virus, a novel virus related to negeviruses, cileviruses and higreviruses. Viruses 6, 4346–4357. doi: 10.3390/v6114346

Kearse, M., Moir, R., Wilson, A., Stones-Havas, S., Cheung, M., Sturrock, S., Buxton, S., Cooper, A., Markowitz, S., Duran, C., Thierer, T., Ashton, B., Meintjes, P., and Drummond, A. (2012). Geneious Basic: An integrated and extendable desktop software platform for the organization and analysis of sequence data. Bioinformatics 28:1647–1649.

Kitajima, E.W., Chagas, C.M., Rodrigues, J.C.V. (2003). *Brevipalpus*-transmitted plant virus and virus-like diseases: cytopathology and some recent cases. Exp Appl Acarol 30: 135–160.

Kondo, H., Fujita, M, Hisano H, Hyodo K, Andika IB and Suzuki N (2020) Virome Analysis of Aphid Populations That Infest the Barley Field: The Discovery of Two Novel Groups of Nege/Kita-Like Viruses and Other Novel RNA Viruses. Front. Microbiol. 11:509. doi: 10.3389/fmicb.2020.00509

Kondo, H., Fujita, M., Telengech, P., Maruyam, K., Hyodo, K., Tassi, A. D., Ochoa, R., Andika, I. B., & Suzuki, N. (2025). Evidence for the replication of a plant rhabdovirus in its arthropod mite vector. Virus research, 351, 199522. 10.1016/j.virusres.2024.199522

Kondo, H., Maeda, T., & Tamada, T. (2003). Orchid fleck virus: *Brevipalpus* californicus mite transmission, biological properties and genome structure. Experimental and Applied Acarology 30(1-3): 215–223. 10.1023/B:APPA.0000006550.88615.10

Koonin, E.V., Dolja, V.V., Krupovic, M., Kuhn, J.H. (2021). Viruses Defined by the Position of the Virosphere within the Replicator Space. Microbiol Mol Biol Rev. 85(4):e0019320. doi: 10.1128/MMBR.00193-20.

Krogh, A., Larsson, B., von Heijne, G., & Sonnhammer, E.L.L. (2001). Predicting transmembrane protein topology with a hidden Markov model: Application to complete genomes. J. Mol. Biol. 305(3), 567–580.

Kubo, K.S., Novelli, V.M., Bastianel, M., Locali-Fabris, E.C., Antonioli-Luizon, R., Machado, M.A., Freitas-Astúa, J. (2011). Detection of Brevipalpus-transmitted viruses in their mite vectors by RT-PCR. Exp Appl Acarol. 54(1):33–9. doi: 10.1007/s10493-011-9425-9.

Kuchibhatla, D.B., Sherman, W.A., Chung BY, Cook S, Schneider G, Eisenhaber B, Karlin DG. (2014). Powerful sequence similarity search methods and in-depth manual analyses can identify remote homologs in many apparently "orphan" viral proteins. J Virol. 88(1):10–20. doi: 10.1128/JVI.02595-13.

Kumar, S., Stecher, G. & Tamura, K. (2016). MEGA 7: Molecular Evolutionary Genetics Analysis version 7.0 for bigger datasets. Mol. Biol. Evol. 33, 1870–1874.

Langmead, B., Salzberg, S. (2012). Fast gapped-read alignment with Bowtie 2. Nat Methods 9, 357–359. 10.1038/nmeth.1923

Larrea-Sarmiento, A., Olmedo-Velarde, A., West-Ortiz, M., Stuehler, D., Hosseinzadeh, S., Coleman, A., Preising, S., Parker, G., Fei, Z., Heck, M. (2024). bioRxiv 2024.08.15.608128; doi: 10.1101/2024.08.15.608128

Lee, H., Lee, C., Tang, J.T., Loh, T.P., Koay, E.S.-C. (2016). Contamination-controlled high-throughput whole genome sequencing for influenza A viruses using the MiSeq sequencer. Sci Rep 6, 33318 10.1038/srep33318

Li, R., Mock, R., Huang, Q., Abad, J., Hartung, J., and Kinard, G. (2008). A reliable and inexpensive method of nucleic acid extraction for the PCR-based detection of diverse plant pathogens. J. Virol. Methods 154, 48–55. doi: 10.1016/j.jviromet.2008.09.008

Lusk, R.W. (2014). Diverse and Widespread Contamination Evident in the Unmapped Depths of High Throughput Sequencing Data. PLoS ONE 9(10): e110808. 10.1371/journal.pone.0110808

Malapi-Wight, M., Adhikari, B., Zhou, J., Hendrickson, L., Maroon-Lango, C.J., McFarland, C., Foster, J.A., Hurtado-Gonzales, O.P. (2021). HTS-Based Diagnostics of Sugarcane Viruses: Seasonal Variation and Its Implications for Accurate Detection. Viruses 13(8):1627. doi: 10.3390/v13081627.

Massart, S., Chiumenti, M., De Jonghe, K., Glover, R., Haegeman, A., Koloniuk, I., Kominek, P., Kreuze, J., … Candresse, T. (2018). Virus detection by high-throughput sequencing of small RNAs: large scale performance testing of sequence analysis strategies. Phytopathology. doi:10.1094/phyto-02-18-0067-r

Melzer, M.J., Sether, D.M., Borth, W.B., Hu, J.S. (2012). Characterization of a virus infecting Citrus volkameriana with citrus leprosis-like symptoms. Phytopathol 102(1): 122–127. doi: 10.1094/PHYTO-01-11-0013.

Melzer, M. J., Simbajon, N., Carillo, J., Borth, W. B., Freitas-Astúa, J., Kitajima, E. W., Neupane, K. R., & Hu, J. S. (2013). A cilevirus infects ornamental hibiscus in Hawaii. Archives of virology 158(11), 2421–2424. 10.1007/s00705-013-1745-0

Menzel, P., Ng, K. & Krogh, A. Fast and sensitive taxonomic classification for metagenomics with Kaiju. Nature Communication 7, 11257 (2016). 10.1038/ncomms11257

Mesa, N. C., Ochoa, R., Welbourn, W. C., Evans, G. A., Moraes, G. J. (2009) A catalog of the Tenuipalpidae (Acari) of the world with a key to genera. Zootaxa. 2098(1):1–185. doi: 10.11646/zootaxa.2098.1.1.

Mironov, S. V., Dabert, J., and Dabert, M. (2012). A new feather mite species of the genus Proctophyllodes Robin, 1877 (Astigmata: Proctophyllodidae) from the long-tailed tit Aegithalos caudatus (Passeriformes Aegithalidae)-morphological description with DNA barcode data. Zootaxa 3253, 54–61.

Navajas, M., Fournier, D., Lagnel, J., Gutierrez, J., and Boursot, P. (1996). Mitochondrial COI sequences in mites: evidence for variations in base composition. Insect Mol. Biol. 5, 281–285. doi: 10.1111/j.1365-2583.1996.tb00102.x

Nunes, M.R.T., Contreras-Gutierrez, M.A., Guzman, H., Martins, L.C., Barbirato, M.F., Savit, C., Balta, V., … Tesh, R.B. (2017). Genetic characterization, molecular epidemiology, and phylogenetic relationships of insect-specific viruses in the taxon Negevirus. Virology 504:152–167. doi: 10.1016/j.virol.2017.01.022.

Nunes, M. A., de Carvalho Mineiro, J. L., Rogerio, L. A., Ferreira, L. M., Tassi, A., Novelli, V. M., Kitajima, E.W., Freitas-Astúa, J. (2018). First Report of *Brevipalpus* papayensis as Vector of Coffee ringspot virus and Citrus leprosis virus C. Plant Dis 102(5): 1046–1046. doi:10.1094/pdis-07-17-1000-pdn

Oksanen J, Simpson G, Blanchet F, Kindt R, Legendre P, Minchin P, O’Hara R, Solymos P, Stevens M, Szoecs E, Wagner H, Barbour M, Bedward M, Bolker B, Borcard D, Borman T, Carvalho G, Chirico M, De Caceres M, Durand S, Evangelista H, FitzJohn R, Friendly M, Furneaux B, Hannigan G, Hill M, Lahti L, Martino C, McGlinn D, Ouellette M, Ribeiro Cunha E, Smith T, Stier A, Ter Braak C, Weedon J (2025). vegan: Community Ecology Package. R package version 2.8–0, https://vegandevs.github.io/vegan/.

Olmedo-Velarde, A., Hu, J., Melzer, M. (2021). A virus infecting Hibiscus-rosa sinensis represents an evolutionary link between cileviruses and higreviruses. Front. Microbiol. 12: 660237.

Olmedo-Velarde, A., Larrea-Sarmiento, A., Wang, X., Hu, J., Melzer, M. (2024). A Breakthrough in Kitavirids: Genetic Variability, Reverse Genetics, Koch’s Postulates, and Transmission of Hibiscus Green Spot Virus 2. Phytopathol. 114:1, 282–293.

Olmedo-Velarde, A., Park, A.C., Sugano, J., Uchida, J.Y., Kawate, M., Borth, W.B., Hu, J.S., Melzer, M.J. (2019) Characterization of Ti Ringspot-Associated Virus, a Novel Emaravirus Associated with an Emerging Ringspot Disease of Cordyline fruticosa. Plant Dis. 103(9):2345–2352. doi: 10.1094/PDIS-09-18-1513-RE.

Pereira, L.R.; Rodrigues, M.C.; Chabi-Jesus, C.; Ramos-González, P.L.; Barbosa, C.J.; Santos, M.G.; Costa, H.; Maro, L.C.; Tassi, A.D.; Kitajima, E.W.;, et al. (2025). Historical and Contemporary Evidence Confirms a Higrevirus as the Causal Agent of Citrus Zonate Chlorosis in Brazil. Viruses 17, 1428. 10.3390/v17111428

Potter, S.C., Luciani, A., Eddy, S.R., Park, Y., Lopez, R. & Finn, R.D. (2018). HMMER web server: 2018 update. Nucleic Acids Res. 46(W1), W200–W204. doi: 10.1093/nar/gky448

Quito-Avila, D.F., Freitas-Astúa, J., and Melzer, M.J. (2021). Bluner-, Cile-, and Higreviruses (Kitaviridae). Encycl. Virol. 2021, 247–251. doi: 10.1016/B978-0-12-809633-8.21248-X

Ramos-González, P.L., Chabi-Jesus, C., Guerra-Peraza, O., Tassi, A.D., Kitajima, E.W., Harakava, R., Salaroli, R.B., Freitas-Astúa, J. (2017) Citrus leprosis virus N: A New Dichorhavirus Causing Citrus Leprosis Disease. Phytopathol 107(8):963–976. doi: 10.1094/PHYTO-02-17-0042-R.

Ramos-González, P.L. Dos Santos, G.F., Chabi-Jesus, C., Harakava, R., Kitajima, E.W., Freitas-Astúa, J. (2020). Passion Fruit Green Spot Virus Genome Harbors a New Orphan ORF and Highlights the Flexibility of the 5′-End of the RNA2 Segment Across Cileviruses. Front. Microbiol. 11:206. doi: 10.3389/fmicb.2020.00206.

Ramos-González, P.L., Pons, T., Chabi-Jesus, C., Arena, G.D. and Freitas-Astua. J. (2021). Poorly Conserved P15 Proteins of Cileviruses Retain Elements of Common Ancestry and Putative Functionality: A Theoretical Assessment on the Evolution of Cilevirus Genomes. Front. Plant Sci. 12:771983. doi: 10.3389/fpls.2021.771983

Rodrigues, J.C.V., Childers, C.C. (2013). *Brevipalpus* mites (Acari: Tenuipalpidae): vectors of invasive, non-systemic cytoplasmic and nuclear viruses in plants. Exp Appl Acarol 59: 165-175.

Roy, A., Choudhary, N., Guillermo, L.M., Shao, J., Govindarajulu, A., Achor, D., Wei, G., Picton, D.D., Levy, L., Nakhla, M.K., Hartung, J.S., Brlansky, R.H. (2013). A novel virus of the genus Cilevirus causing symptoms similar to citrus leprosis. Phytopathol 103(5):488–500. doi: 10.1094/PHYTO-07-12-0177-R. PMID: 23268581.

Roy, A., Hartung, J. S., Schneider, W. L., Shao, J., Leon, M. G., Melzer, M. J., Beard, J. J., Otero-Colina, G., Bauchan, G. R., Ochoa, R., and Brlansky, R. H. (2015) Role bending: Complex relationships between viruses, hosts, and vectors related to citrus leprosis, an emerging disease. Phytopathol 105:1013–1025.

Sameroff, S., Tokarz, R., Charles, R.A. Jain, K., Oleynik, A., Che, X., Georges, K., Carrington, C.V., Lipkin, W.I., Oura, C. (2019). Viral Diversity of Tick Species Parasitizing Cattle and Dogs in Trinidad and Tobago. Sci Rep 9, 10421. 10.1038/s41598-019-46914-1

Shi, M., Lin, XD., Tian, JH. et al. (2016). Redefining the invertebrate RNA virosphere. Nature 540, 539–543. 10.1038/nature20167

Soltani, N.; Stevens, K.A.; Klaassen, V.; Hwang, M.-S.; Golino, D.A.; Al Rwahnih, M. (2021) Quality Assessment and Validation of High-Throughput Sequencing for Grapevine Virus Diagnostics. Viruses 13, 1130. 10.3390/v13061130

Sonnenberg, R., Wolte, A. W., and Tautz, D. (2007). An evaluation of LSU rDNA D1-D2 sequences for their use in species identification. Front. Zool. 4:6. doi: 10.1186/1742-9994-4-6

Talavera, G., and Castresana, J. (2007). Improvement of phylogenies after removing divergent and ambiguously aligned blocks from protein sequence alignments. Syst. Biol. 56, 564–577. doi: 10.1080/10635150701472164

Tassi, A.D. (2018). Diversidade morfológica e genética de diferentes espécies de Brevipalpus (Acari: Tenuipalpidae) e suas competencias como vetores de virus. 262. Available at: https://teses.usp.br/teses/disponiveis/11/11135/tde-17072018-160552/publico/Aline_Daniele_Tassi_versao_revisada.pdf

Tassi A. D., Ramos-González, P. L., Flechtmann, C. H. W., Amrine, J. W. Jr, Sarkhosh, A., Freitas-Astua, J., Kitajima, E. W., Marques, J. P. R., Harmon, P. F., Carrillo, D. (2025) Eriophyid Mites Vector the Kitavirus Blueberry Necrotic Ring Blotch Virus: Insights into the Viral Transmission and Its Infection on Blueberry Plants. Phytopathology 115(8):1038–1050. doi: 10.1094/PHYTO-02-25-0063-R

Tassi, A. D., Ramos-González, P. L., Sinico, T. E., Kitajima, E. W., & Freitas-Astúa, J. (2022). Circulative Transmission of Cileviruses in Brevipalpus Mites May Involve the Paracellular Movement of Virions. Frontiers in microbiology, 13, 836743. 10.3389/fmicb.2022.836743

Thompson, J.D., Higgins, D.G., and Gibson, T.J. 1994. CLUSTAL W: improving the sensitivity of progressive multiple sequence alignment through sequence weighting, position-specific gap penalties and weight matrix choice. Nucleic Acids Res. 22:4673–4680.

Untergasser, A., Cutcutache, I., Koressaar, T., Ye, J., Faircloth, B.C., Remm, M., et al. (2012). Primer3 - new capabilities and interfaces. Nucleic Acids Res. 40(15): e115

Valles S.M., Chen, Y., Firth, A.E., Guerin, D.M.A., Hashimoto, Y., Herrero, S., de Miranda, J.R., Ryabov, E. (2017). ICTV Virus Taxonomy Profile: Dicistroviridae. J Gen Virol. 98:355–6.

van Oers, M.M. (2010). Genomics and Biology of Iflaviruses. In K. Johnson, & S.. Agari (Eds.), Insect Virology (pp. 231–250). Caister Academic Press.

Vasilakis, N., Forrester, N.L., Palacios, G., Nasar, F., Savji, N., Rossi, S.L., Guzman, H., Wood, T.G., … Tesh, R. B. (2013). Negevirus: a proposed new taxon of insect-specific viruses with wide geographic distribution. Journal of virology, 87(5), 2475–2488. 10.1128/JVI.00776-12

Villamor, D.E.V., Ho, T., Al Rwahnih, M., Martin, R.R., Tzanetakis, I.E. (2019). High Throughput Sequencing For Plant Virus Detection and Discovery. Phytopathol. 109(5): 716–725. 10.1094/PHYTO-07-18-0257-RVW

Whitfield, A.E., Huot, O.B., Martin, K.M., Kondo, H., Dietzgen, R.G. (2018). Plant rhabdoviruses—their origins and vector interactions. Current Opinion in Virology 33: 198–207.

Zhang, Y. Z., Shi, M., & Holmes, E. C. (2018). Using Metagenomics to Characterize an Expanding Virosphere. Cell, 172(6), 1168–1172. 10.1016/j.cell.2018.02.043

