## Supplementary for "Viromics in flat mites from Hawaii shows abundant arrays of viruses, expands the evolutionary origin of plant viruses, and provides a surveillance tool for *Brevipalpus*-transmitted viruses"

**Supplementary Material**

**Table S1.** BLASTn searches results of the partial 28S rRNA and COI sequences amplified from the mite samples used in this study.

| **Mite** | **Host** | **Location^3^** | **28S rRNA** | | | **Cytochrome oxidase I (COI)** | | | |
| --- | --- | --- | --- | --- | --- | --- | --- | --- | --- |
| **Sample** |  |  | **Nucleotide** | **GenBank** | **Query** | | **Nucleotide** | **GenBank** | **Query** |
| **ID** |  |  | **identity** | **Accession** | **coverage** | | **identity** | **Accession** | **coverage** |
| Citrus O1 | *Citrus* spp. | Waimanalo | *B. azores* 100% | [MK919272](https://www.ncbi.nlm.nih.gov/nucleotide/MK919272.1?report=genbank&log$=nucltop&blast_rank=1&RID=R447B8C801N) | 96% | | *B. azores* 99.72% | [MK499460](https://www.ncbi.nlm.nih.gov/nucleotide/MK499460.1?report=genbank&log$=nucltop&blast_rank=1&RID=R43NHJWY013) | 87% |
| Citrus O2 | *Citrus* spp. | Pearl City | *B. yothersi* 99.66% | MK293683 | 96% | | *B. yothersi* 99.4% | KX100285 | 85% |
| Citrus O3 | *Citrus* | Manoa | *B. azores* 98.86% | MK919272 | 92% | | *B. yothersi* 99.71% | KF954967 | 90% |
|  | *reticulata* |  | *B. yothersi* 99.66% | MK293630 | 93% | |  |  |  |
| Citrus H1 | *Citrus* spp. | Waiakea | *B. yothersi* 100% | MK293678 | 93% | | *B. yothersi* 99.71% | KF954967 | 84% |
| Hibiscus O1 | *Hibiscus arnottianus* | Manoa | *B. papayensis* 99.33% | [MT664800](https://www.ncbi.nlm.nih.gov/nucleotide/MK919272.1?report=genbank&log$=nucltop&blast_rank=1&RID=R447B8C801N) | 94% | | *B. papayensis* 98.55% | [KF954950](https://www.ncbi.nlm.nih.gov/nucleotide/MK919272.1?report=genbank&log$=nucltop&blast_rank=1&RID=R447B8C801N) | 84% |
| Hibiscus O2 | *Hibiscus arnottianus* | Manoa | *B. papayensis* 99.33% | [MT664800](https://www.ncbi.nlm.nih.gov/nucleotide/MK919272.1?report=genbank&log$=nucltop&blast_rank=1&RID=R447B8C801N) | 94% | | *B. papayensis* 99.02% | [KF954950](https://www.ncbi.nlm.nih.gov/nucleotide/MK919272.1?report=genbank&log$=nucltop&blast_rank=1&RID=R447B8C801N) | 82% |
| Hibiscus O3 | *Hibiscus rosa-sinensis* | Pearl City | *B. obovatus* 100% | [MK293694](https://www.ncbi.nlm.nih.gov/nucleotide/MK919272.1?report=genbank&log$=nucltop&blast_rank=1&RID=R447B8C801N) | 90% | | *B. obovatus* 100% | [MK499465](https://www.ncbi.nlm.nih.gov/nucleotide/MK919272.1?report=genbank&log$=nucltop&blast_rank=1&RID=R447B8C801N) | 86% |
| Passionfruit O1 | *Passiflora edulis* | Ala Wai | *B. yothersi* 100% | MK293630 | 94% | | *B. yothersi* 99.74% | MH605068 | 100% |
| Passionfruit O2 | *Passiflora edulis* | Manoa | *B. yothersi* 100% | MK293630 | 93% | | *B. yothersi* 99.71% | KF954967 | 89% |
| Passionfruit O3 | *Passiflora edulis* | Waimanalo | *B. yothersi* 100% | MK293630 | 93% | | *B. yothersi* 99.71% | KF954967 | 89% |
| Papaya O1 | *Carica papaya* | Poamoho | *B. papayensis* 99.33% | [MT664800](https://www.ncbi.nlm.nih.gov/nucleotide/MK919272.1?report=genbank&log$=nucltop&blast_rank=1&RID=R447B8C801N) | 94% | | *B. papayensis* 98.84% | [KF954950](https://www.ncbi.nlm.nih.gov/nucleotide/MK919272.1?report=genbank&log$=nucltop&blast_rank=1&RID=R447B8C801N) | 89% |
| Papaya O2 | *Carica papaya* | Manoa | *B. yothersi* 99.52% | MT812697 | 100% | | *B. yothersi* 99.71% | KF954967 | 89% |
| Coffee H1 | *Coffea* sp. | Kona | *B. papayensis* 99.33% | [MT664800](https://www.ncbi.nlm.nih.gov/nucleotide/MK919272.1?report=genbank&log$=nucltop&blast_rank=1&RID=R447B8C801N) | 94% | | *B. papayensis* 98.84% | [KF954950](https://www.ncbi.nlm.nih.gov/nucleotide/MK919272.1?report=genbank&log$=nucltop&blast_rank=1&RID=R447B8C801N) | 84% |
| Coffee O1 | *Coffea* sp. | Poamoho | *B. papayensis* 98.49-99.16% | [MT664798 MT664800](https://www.ncbi.nlm.nih.gov/nucleotide/MK919272.1?report=genbank&log$=nucltop&blast_rank=1&RID=R447B8C801N) | 94% | | *B. papayensis* 98.84% | [KF954950](https://www.ncbi.nlm.nih.gov/nucleotide/MK919272.1?report=genbank&log$=nucltop&blast_rank=1&RID=R447B8C801N) | 89% |
| Pineapple O1 | *Ananas comosus* | Poamoho | *Raioella indica* 78.62% | JF928445 | 93% | | *Dolichotetranychus* sp.  *99.73%* | MH606191 | 95% |

**Table S2.** Primer sequences used in RT-PCR assays for virus presence confirmation from the different mite samples used in this study.

| **Virome** | **Virus contig name** | **Forward (5' - 3')** | **Reverse (5' - 3')** | **Product size (bp)** |
| --- | --- | --- | --- | --- |
| *Brevipalpus*- | Brevipalpus-associated ourmiavirus 1 | CATCAACGCCCTGGAGTACC | CCTAATTGCCGGCCTTCACT | 551 |
| associated | Brevipalpus-associated narnavirus 1 | CCCATCGGACTTCCTCTCCT | AGTCTCCCTCCGTTGCCTAA | 251 |
| virome | Brevipalpus-associated picornavirus 1 | CAATGGAGTGCGTGGATGGA | ATGTCTCTCGGGCTCACAGA | 656 |
| (citrus) | Brevipalpus-associated picornavirus 2 | TGACACGGAAGAGCAGAGGT | TTCGTGGTCATTTGCGTGCA | 566 |
|  | Brevipalpus-associated picornavirus 3 | CGCTGTTGGACACCGTAGTT | GGGCACGAAAGGTTCCTGAT | 427 |
|  | Brevipalpus-associated picornavirus 4 | AGATCCTTCCGAGCCACCTT | CAGGAGACAGTCGAGTTGGC | 246 |
|  | Brevipalpus-associated picornavirus 5 | ACAGGAAATGGCAAAGGTGTC | ACTGCTGCATCGACAAATTGG | 176 |
|  | Brevipalpus-associated picornavirus 6 | ACTCCGATACAATCATTGCAGGA | GTACGTGGGAGTTGGTTCGG | 243 |
|  | Brevipalpus-associated tombusvirus 1 | TGCGCGAACCTCTTTATGCT | TCACATCAAACGCCAGCTCA | 255 |
|  | Brevipalpus-associated tombusvirus 2 | CCAACCATGTCCCGTCAACA | CGATTTCAACACCGGCCTTG | 353 |
| *Brevipalpus*- | Brevipalpus-associated bluner-like virus RNA 1 | GGTTGATGCATGGTGAGGGT | CCCAACGCCCTAACCAACAT | 430 |
| associated | Brevipalpus-associated bluner-like virus RNA 2 | TCATGGTTGGCAGGCTTTGT | TTCCACAGTCTCTCGCAACG | 409 |
| Virome | Brevipalpus-associated negevirus 1 | ACCCTAGATGTGTCCTCGGT | GCGTCACATCATCGTAGGCT | 511 |
| (hibiscus) | Brevipalpus-associated ourmiavirus 2 | AACGCGTTCCTTCGACTCTC | CTAAGCGACCGTCCTTCCAC | 666 |
|  | Brevipalpus-associated narnavirus 2 | TAAACGGTTGGGCAGCAGAG | GCTCTTCGGGTGCCTTATCG | 213 |
|  | Brevipalpus-associated ourmiavirus 3 | GTACTGGGCAAGCTCAAGGT | AGCAAGAGATCATCGCCGTT | 200 |
|  | Brevipalpus-associated ourmiavirus 4 | CAAGCGTTTGGGACCTAGCA | AAGCGCTCATTCACAAGGGT | 292 |
|  | Brevipalpus-associated narnavirus 3 | GCGGGATTGACGGTTACCTC | AAGACTCGCGAATCCAGCTC | 321 |
|  | Brevipalpus-associated ourmiavirus 5 | GGTCAACTCATGGGTTCGCT | TTTCTTGGGTGCCTGTGTCC | 675 |
|  | Brevipalpus-associated picornavirus 7 | CCCTGAACGCCGAATCCATT | ATGTGTGGGCTGAGGTCATG | 329 |
|  | Brevipalpus-associated picornavirus 8 | GGTAGTCCCTCAGGTAGTGCT | GAGCCTGAACGTCGAATCCA | 398 |
|  | Brevipalpus-associated picornavirus 9 | CCTAGCTGCAGACGGGAAAG | CATGGCAGAGAAACCGGACA | 470 |
|  | Brevipalpus-associated picornavirus 10 | TCTGGGAGTGACTTGTGGACT | GCCACTCCAACTCCACATTTG | 405 |
|  | Brevipalpus-associated picornavirus 11 | AGTCGCAGGTGTAGCATTCG | GGACAATAACATTCCCGAAGTGC | 226 |
|  | Brevipalpus-associated tombusvirus 3 | CCCTCAAGTTCGGCCATGAA | AGGCACAAATCCCTCCGTTG | 293 |
|  | Hibiscus green spot virus 2 RNA 1 | TGTATGGTGCCCGTGTTGTC | CGTCACACCACTCAGCAACA | 232 |
|  | Hibiscus green spot virus 2 RNA 2 | GCTGAACGGTGTTCCTGGTG | CCCGTGCATGAACAAGTCGA | 450 |
|  | Hibiscus green spot virus 2 RNA 3 | CTCGGTGCTCTTGTTGTTGC | GCAAAGACACGAACCCAAGC | 526 |
| *Brevipalpus*- | Brevipalpus-associated picornavirus 12 | TACGAGGATGATGGCGGTGA | TGCCGGACGAGGGTAATCTA | 267 |
| associated | Brevipalpus-associated picornavirus 13 | AGTTAGTTTACCCTCGCGTGTT | AGAGGCCTCACGTTGCTCTA | 160 |
| virome | Citrus leprosis virus C2 RNA 1 | ACAAGATGGCGGACGAACTG | AGCCATGTCATCGGGATCCA | 386 |
| (papaya) | Citrus leprosis virus C2 RNA 2 | GATCACCGTAGTTTCATGCG | CCAACGCGGTTAACAACTC | 129 |
|  | Papaya ringspot virus | TGACGGCTGGGTGTACTGT | TGCCGCTGTGATTGCCTC | 596 |
| *Brevipalpus*- | Citrus leprosis virus C2 RNA 1 | ACAAGATGGCGGACGAACTG | AGCCATGTCATCGGGATCCA | 386 |
| associated virome | Citrus leprosis virus C2 RNA 2 | GATCACCGTAGTTTCATGCG | CCAACGCGGTTAACAACTC | 129 |
| (passionfruit) | Brevipalpus-associated reovirus | CGGCCGGATTAACTGAAGAACA | GGATGTCCTGTAGCCATTAATCCT | 358 |
| Tenuipalpid-associated | Dolichotetranychus-associated negevirus contig | AGCCGTGATTCAATGCGAGA | ATAACGCCGACACCCACTTC | 266 |
| virome | Dolichotetranychus-associated solemovirus | ATGTTGTCTGGGCGGATCAC | TTAAGTTCCGGCTGCCTGAC | 545 |
| (pineapple-coffee) | Dolichotetranychus-associated cile-like virus contig | ACTCCGTTGGTTGTGGGTTC | CCCTTCCAACCAATCAGCCA | 465 |

**Table S3.** Alpha-diversity metrics of each virome pool. Richness (number of viral taxonomic groups), Shannon diversity, and Pielou’s evenness were calculated separately for the count and read-abundance datasets, each aggregated by assigned viral taxonomic group.

| **Pool** | **Richness** | **Count data** | | **Read abundance data** | |
| --- | --- | --- | --- | --- | --- |
|  |  | **Shannon** | **Evenness** | **Shannon** | **Evenness** |
| Citrus | 4 | 1.09 | 0.79 | 0.19 | 0.14 |
| Hibiscus | 6 | 1.62 | 0.90 | 1.41 | 0.79 |
| Passionfruit | 2 | 0.69 | 1.00 | 0.49 | 0.70 |
| Papaya | 3 | 1.04 | 0.95 | 0.34 | 0.31 |
| Pineapple | 3 | 1.10 | 1.00 | 1.09 | 0.99 |

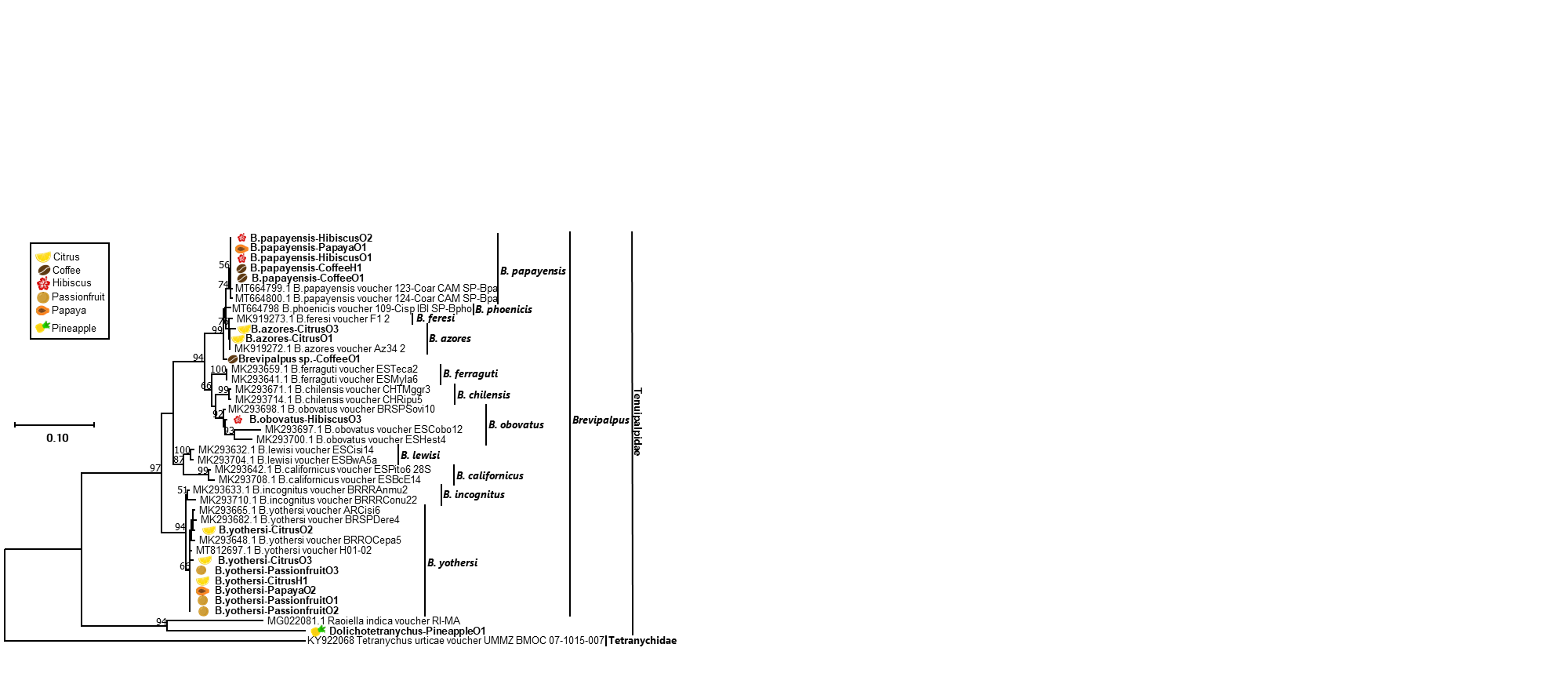

**Figure S1. Phylogenetic relationships using partial 28S rRNA gene sequences from several tenuipalpid mites classified within *Brevipalpus*, *Raioella* and *Dolichotetranychus*.** Relationships were based on a multiple nucleotide sequence alignment using CLUSTAL and inferred using the Maximum Likelihood algorithm implemented in MEGA 7.0.25. Bootstrap values greater than 50 are shown above the branches after 1,000 repetitions. No *Dolichotetranychus* 28S rRNA sequence was found in the GenBank database. A homolog partial 28S rRNA gene sequence from *Tetranychus urticae* (Tetranychidae) was used as an outgroup. The scale at the left indicates the number of substitutions per given branch length. Tenuipalpid mite samples from in this study are bold and placed next to the respective icon of the plant/fruit from which they were collected.

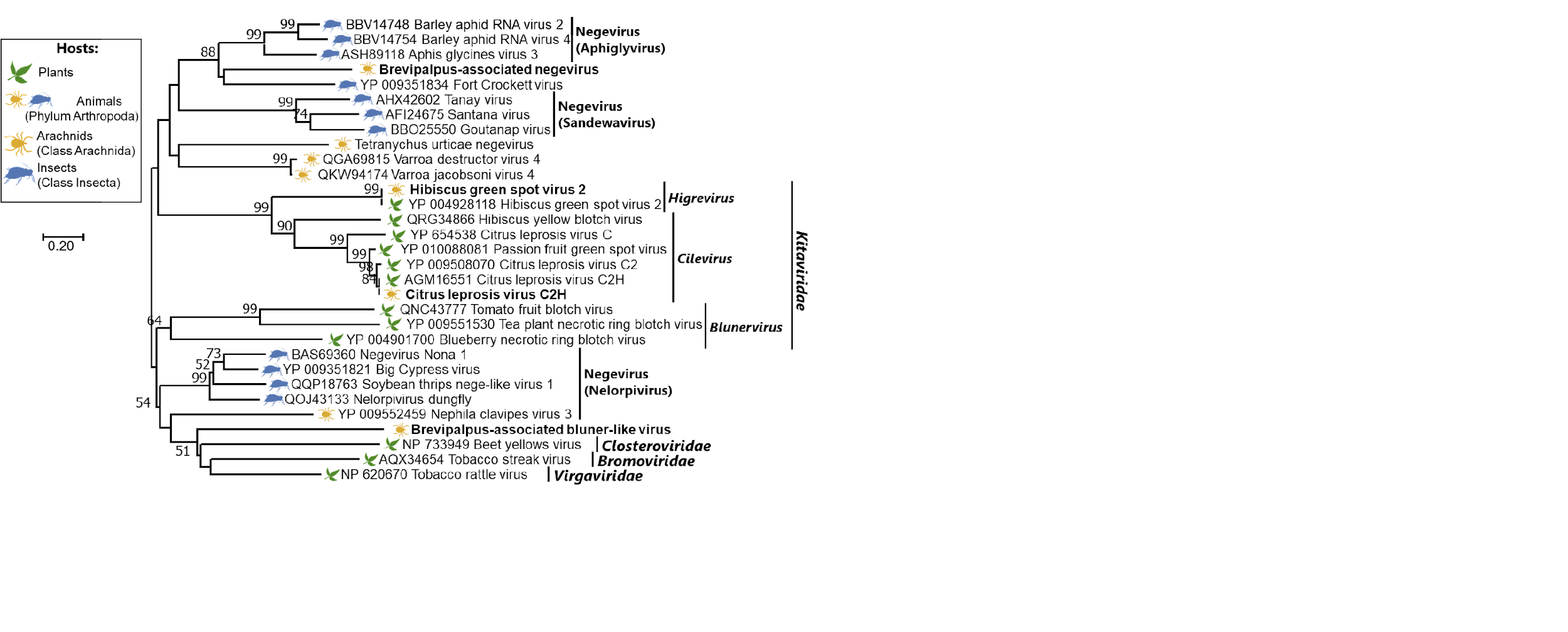

**Figure S2. Phylogenetic relationships inferred using helicase (HEL) conserved domain found in the contig sequences of the putative new negeviruses and kitavirids in tenuipalpid mite samples.** The specific mite sample that each putative new virus was detected from is detailed in Table 3. Brevipalpus-associated bluner-like virus corresponds to the RdRp domain present in the Brevipalpus-associated bluner-like virus contig 1 (RNA 1) (Figure 3).

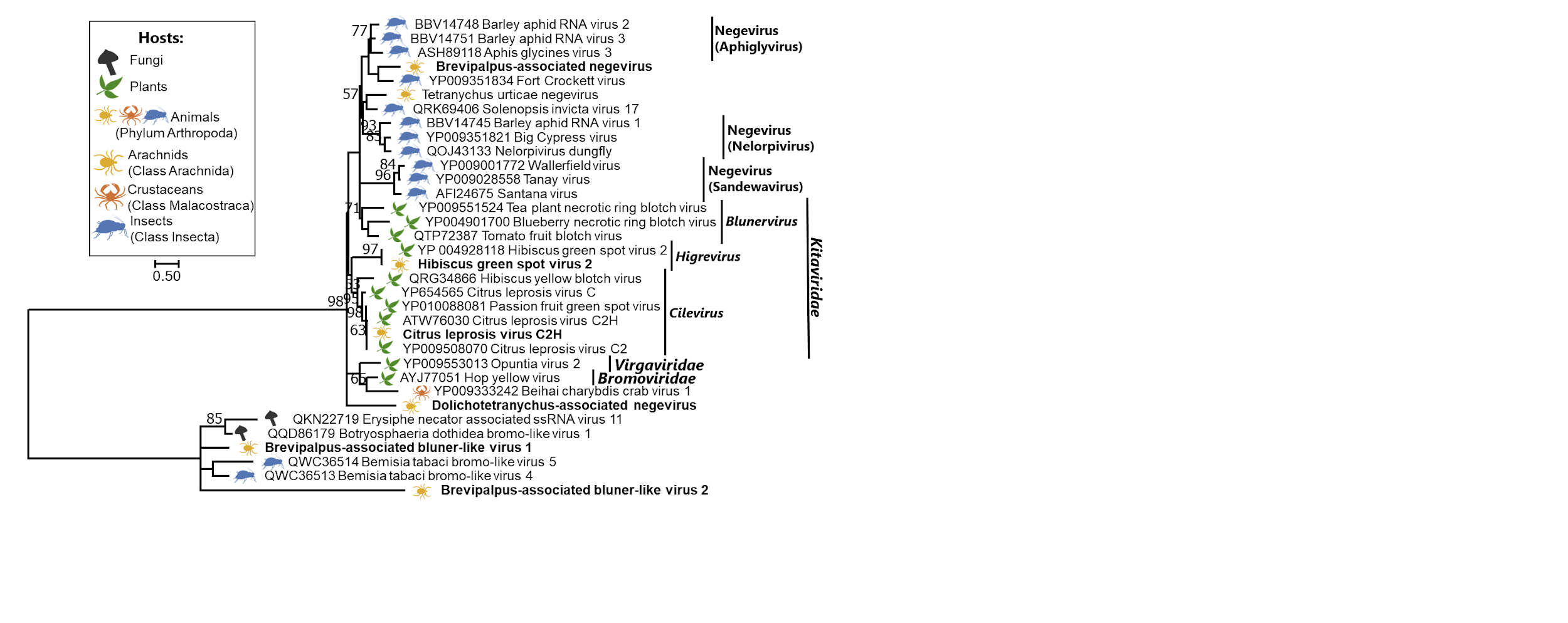

**Figure S3. Phylogenetic relationships inferred using the methyltransferase (MET) conserved domain found in the contig sequences of the putative new negeviruses and kitavirids in tenuipalpid mite samples.** The specific mite sample that each putative new virus was detected from is detailed in Table 3. Brevipalpus-associated bluner-like virus 1 and 2 correspond to the MET domains present in the Brevipalpus-associated bluner-like virus contig 1 (RNA 1) and 2 (RNA 2), respectively (Figure 3).

0.67

0

0.14

0.17

0.29

0.25

0

0.29

0.25

0.2

0.22

0

0.02

0

0.04

0.36

0.07

0.11

0

0.03

Pineapple

Papaya

Passionfruit

Hibiscus

Citrus

Citrus

Hibiscus

Passionfruit

Papaya

Pineapple

Jaccard similarity

0.00

0.25

0.50

0.75

1.00

Bray–Curtis similarity

0.00

0.25

0.50

0.75

1.00

**Figure S4.** Heatmap showing pairwise β-diversity among virome pools based on Jaccard (blue) and Bray–Curtis (red) similarity metrics. The lower triangle represents Jaccard similarity calculated from the count dataset, and the upper triangle represents Bray–Curtis similarity calculated from the read-abundance dataset. Similarity values were derived as 1 - dissimilarity, where 0 = completely dissimilar and 1 = identical.
